# The *C. elegans* Anchor Cell Transcriptome: Ribosome Biogenesis Drives Cell Invasion through Basement Membrane

**DOI:** 10.1101/2022.12.28.522136

**Authors:** Daniel S. Costa, Isabel W. Kenny-Ganzert, Qiuyi Chi, Kieop Park, Laura C. Kelley, Aastha Garde, David Q. Matus, Junhyun Park, Shaul Yogev, Bob Goldstein, Theresa V. Gibney, Ariel M. Pani, David R. Sherwood

**Affiliations:** Department of Biology, Duke University, Box 90338, Durham, NC 27708, USA; Department of Cell Biology, Duke University Medical Center, Durham, NC 27708, USA; Department of Molecular Biology, Princeton University, Princeton, NJ 08544, USA; Howard Hughes Medical Institute, Princeton University, Princeton, NJ 08544, USA; Department of Biochemistry and Cell Biology, Stony Brook University, Stony Brook, NY 11794, USA; Department of Neuroscience, Yale School of Medicine, New Haven, CT 06510, USA; Department of Biology, University of North Carolina at Chapel Hill, Chapel Hill, NC 27599, USA.; Department of Biology, University of Virginia, Charlottesville, VA 29903, USA.; Department of Cell Biology, University of Virginia School of Medicine, Charlottesville, VA 29904, USA

**Keywords:** cell invasion, transcriptome, basement membrane, ribosome biogenesis, endomembrane expansion, Translationally Controlled Tumor Protein (TCTP)

## Abstract

Cell invasion through basement membrane (BM) barriers is important in development, immune function, and cancer progression. As invasion through BM is often stochastic, capturing gene expression profiles of cells actively transmigrating BM *in vivo* remains elusive. Using the stereotyped timing of *C. elegans* anchor cell (AC) invasion, we generated an AC transcriptome during BM breaching. Through a focused RNAi screen of transcriptionally enriched genes, we identified new invasion regulators, including TCTP (Translationally Controlled Tumor Protein). We also discovered gene enrichment of ribosomal proteins. AC-specific RNAi, endogenous ribosome labeling, and ribosome biogenesis analysis revealed a burst of ribosome production occurs shortly after AC specification, which drives the translation of proteins mediating BM removal. Ribosomes also strongly localize to the AC’s endoplasmic reticulum (ER) and the endomembrane system expands prior to invasion. We show that AC invasion is sensitive to ER stress, indicating a heightened requirement for translation of ER trafficked proteins. These studies reveal key roles for ribosome biogenesis and endomembrane expansion in cell invasion through BM and establish the AC transcriptome as a resource to identify mechanisms underlying BM transmigration.

## INTRODUCTION

Basement membrane (BM) is a thin, dense, laminin and type IV collagen rich extracellular matrix (ECM) that enwraps most tissues (Yurchenco, 2011). BM mechanically supports tissues and has formidable barrier properties, which prevents cell movement between tissues (Bunt et al., 2010; Jayadev et al., 2022; Sherwood, 2021; Yurchenco, 2011). Yet, a number of cells acquire the ability to breach BM to exit or enter tissues. For example, vertebrate neural crest cells, gastrulating mesoderm and endoderm precursor cells, and limb muscle cell precursor cells undergo an epithelial-to-mesenchymal transition (EMT) and invade through the BM to migrate away from the epithelial tissue (Gros and Tabin, 2014; Leonard and Taneyhill, 2020; Nakaya et al., 2008). Macrophages and neutrophils also breach BM to enter tissues during immune cell survellience (Bahr et al., 2022; van den Berg et al., 2019). Cell invasive behavior is misregulated in developmental disorders, inflammatory diseases, and cancer (Fisher, 2015; Novikov et al., 2021; Xing et al., 2016). Thus, understanding the mechanisms underlying cell invasion is of basic biological and clinical importance.

Studies examining BM transmigrating cells have revealed that invasive cells use specialized cellular protrusions to penetrate BM, which harbor over 300 different proteins, such as adhesion receptors, cytoskeletal regulators, proteases, and signaling proteins (Cambi and Chavrier, 2021; Ezzoukhry et al., 2018; Linder et al., 2022). In addition, BM transmigrating cells have robust metabolic networks that fuel the invasive cytoskeletal and adhesion machinery (Garde et al., 2022; Garde and Sherwood, 2021; Papalazarou et al., 2020). Together, these specialized features require the transcription of numerous genes controlled by pro-invasive transcription factor networks (Medwig-Kinney et al., 2020; Pastushenko and Blanpain, 2019). How invasive cells translate the mRNAs encoding numerous new proteins required for invasion is less understood. Notably, recent studies have revealed that ribosome biogenesis is required for the execution of EMT in multiple cancer cell lines and neural crest cells during development (Prakash et al., 2019). Whether ribosome biogenesis promotes BM invasion during EMT or is involved in another aspect of the EMT program is unknown.

Anchor cell (AC) invasion in *Caenorhabditis elegans* is a visually and genetically tractable model of BM transmigration (Sherwood and Plastino, 2018). The AC is a specialized uterine cell that invades through the juxtaposed uterine and vulval BM to initiate uterine-vulval connection during larval development. The AC shares many similarities with other invasive cells, including pro-invasive transcription factor networks, specialized invasive protrusions, and a complex metabolic system (Garde et al., 2022; Hagedorn et al., 2013; Kelley et al., 2019; Medwig-Kinney et al., 2020; Naegeli et al., 2017; Sherwood et al., 2005). Complete disruption of AC invasion perturbs uterine-vulval attachment and leads to a protruded vulva (Pvl) phenotype and many RNAi and genetic screens have used this phenotype to identify invasion promoting genes (Kelley et al., 2019; Lattmann et al., 2022; Lohmer et al., 2016; Matus et al., 2010; Matus et al., 2015; Sherwood et al., 2005). Genes encoding proteins required for viability, as well as subtle invasion regulators, however, have likely been missed in Pvl screens.

Here, we have taken advantage of the stereotyped timing of AC invasion and established the AC gene expression profile/transcriptome during BM breaching to gain deeper insight into BM invasion. We identified ∼1,500 genes enriched in the AC during BM breaching and conducted a focused RNAi screen on a subset of these and discovered new invasion regulators, including TCT-1, an ortholog of the vertebrate Translationally Controlled Tumor Protein (TCTP). The AC transcriptome also revealed enrichment of genes encoding ribosomal proteins. Through AC-specific RNAi knockdown, endogenous ribosomal protein labeling, and ribosome biogenesis analysis, we discovered that a burst of ribosome biogenesis occurs shortly after AC specification and promotes BM invasion by driving the translation of numerous pro-invasive proteins, including a heavy reliance on proteins trafficked through the endomembrane system. Together, our studies uncover the transcriptome of an invasive cell during BM breaching and reveal a key role for ribosome biogenesis in faciliating BM invasion.

## RESULTS

### The generation of RNA-seq libraries of invading *C. elegans* anchor cells (ACs)

AC invasion through BM is highly stereotyped and occurs over a 90-minute period during the L3 larval stage. The time course of invasion can be staged by the divisions of the underlying P6.p vulval precursor cell (VPC), as invasion occurs in coordination with P6.p VPC divisions (Sherwood and Sternberg, 2003). The AC is specified during the late L2-to-early L3 stage (early P6.p 1-cell stage, ∼ 6 hours before invasion (Kimble, 1981; Ziel et al., 2009)), and AC invasion initiates at the P6.p 2-cell stage with protrusions that breach and clear the BM by the P6.p 4-cell stage (Fig. 1A) (Hagedorn et al., 2013).

**Figure 1.**
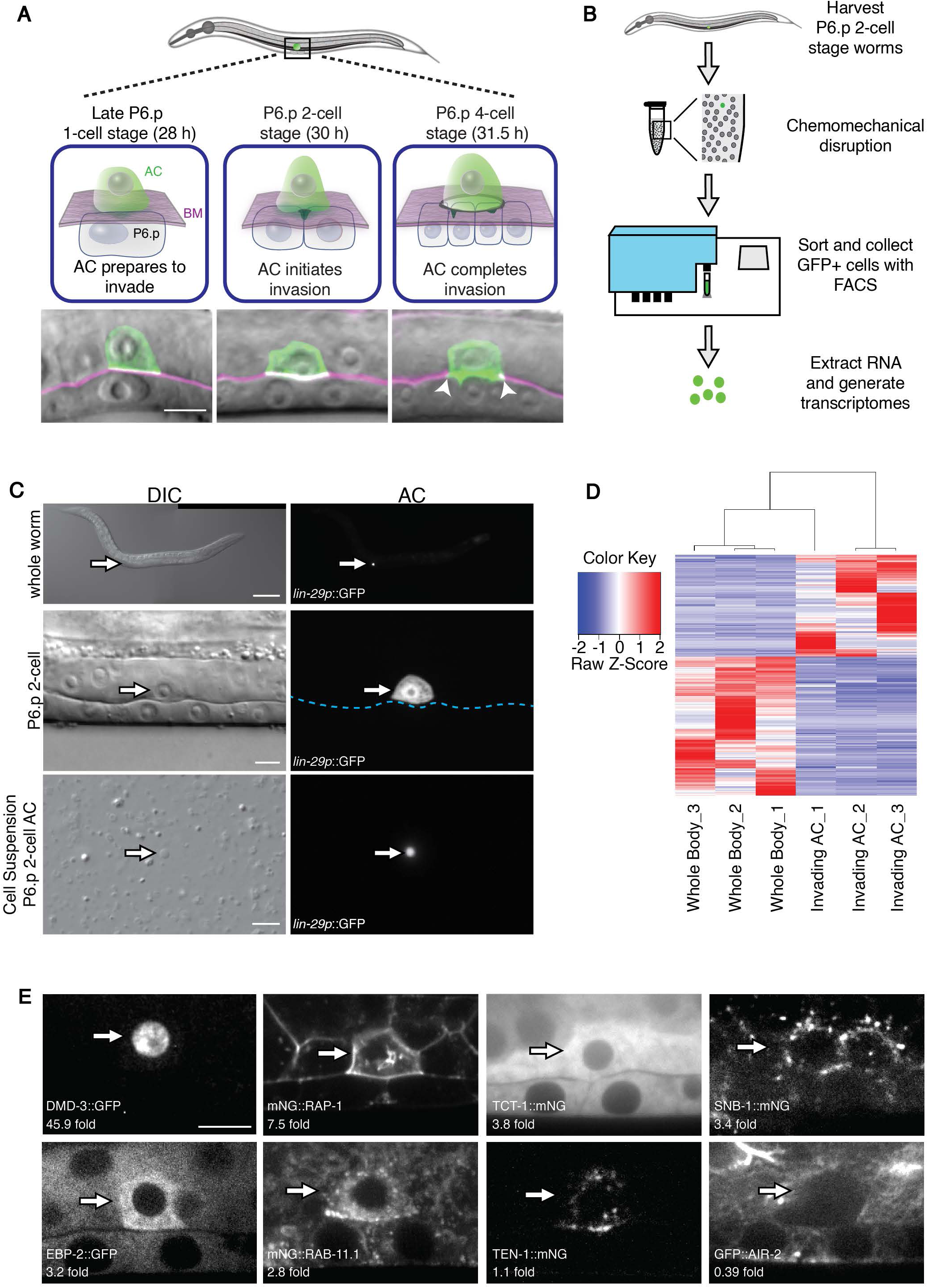
Creation, analysis and validation of the anchor cell (AC) transcriptome. (A) *C. elegans* AC (green) invasion through BM (magenta) from the 1° vulval precursor cell (VPC) late P6.p 1-cell stage to the P6.p 4-cell stage. The top shows a schematic lateral view of invasion and the bottom shows micrographs of the AC (plasma membrane labeled by *cdh-3p*::mCherry::PLCδ^PH^) and BM (viewed with *lam-1p*::LAM-1::GFP) fluorescence overlaid on differential interference contrast (DIC) images. Arrowheads mark the BM breach. Timeline shown is in hours post-hatching at 20°C. (B) Outline of cell isolation, FACS, and generation of transcriptomes. (C) Animals at the time of AC isolation (left, DIC; right, *lin-29p*::GFP AC fluorescence, dotted blue line represents BM). AC in the intact animal (arrows, top, 100X magnification; middle 1000X magnification) and in cell suspension (arrows, bottom). (D) Hierarchical clustering of genes with a significant p-value (≤0.05). Raw Z-score values for the top 200 differentially expressed genes shown (red = high, blue = low). (E) Endogenously tagged proteins expressed in the AC (arrows) and fold change of AC mRNA levels compared to the whole-body (WB) gene expression. Abbreviations: Basement membrane (BM), anchor cell (AC), hours (h). Scale bars: (C) top scale bar, 40 µm. All other scale bars, 5 µm.

To complement previous phenotypic screening approaches used to identify genes that promote cell invasion, we sought to characterize the gene expression profile of the AC at the time of BM breaching. We used synchronized worm culturing, single-cell isolation techniques, and FACS (Fluorescence-Activated-Cell-Sorting) to isolate ACs during BM breaching (Fig. 1B) (Spencer et al., 2014; Zhang et al., 2011). For fluorescent identification of ACs, we found that the *lin-29* promoter (*lin-29p::*GFP) (Garde et al., 2022) drove strong and specific GFP expression in the AC during BM invasion (Fig. 1C). Chemomechanical disruption of *C. elegans* at the mid-L3 P6.p 2-cell stage resulted in a cell suspension with rare highly fluorescent ACs (Fig. 1C). We isolated ACs using FACS followed by RNA-sequencing (RNA-seq) to generate three AC libraries (between ∼80,000-to-150,000 ACs/library, 8-10 ng of total RNA). We also generated three whole-body (WB) gene expression profiles at the same mid-L3 stage. Gene body coverage showed unbiased representation of the 5’ and 3’ ends of transcripts in all of the WB libraries and 2 of 3 AC libraries (Fig. S1). One AC library had a slight 3’ bias, likely because of the use of poly(dT) for cDNA synthesis. Hierarchical clustering of all genes with a significant p-value (≤0.05) showed that specific genes differentiate the AC and WB libraries (Fig. 1D). Using the three AC libraries, we created an AC transcriptome (Table S1), and detected 12,682 genes with at least 10 reads in one of the AC libraries. Comparing the gene expression in the AC dataset versus WB dataset, we identified 1,502 transcripts with significantly elevated expression (2-fold higher, Benjamini–Hochberg adjusted P-value < 0.1) (Table S2).

To validate our AC transcriptome, we compiled a list of 52 genes previously shown with fluorescent reporters to be expressed in the AC during invasion (Table S3) and found that 51/52 of these genes were present with at least 10 copies in one of the AC libraries (Tables S1,S3). We also created genome edited *C. elegans* strains where mNeonGreen (mNG) was inserted in frame with the protein coding region of five highly enriched genes (2-fold or greater) —*tct-1* (translationally controlled tumor protein), s*nb-1* (synaptobrevin), *eif-1.A* (translation initiation factor EIF1A), *lin-3* (EGF ligand) and *rab-11.1* (Rab11 small GTPase) and examined 16 other previously tagged genes that were enriched at various levels in the AC (Tables S3,S4). Examination of ACs at the time of BM breaching (P6.p 2-cell stage, n ≥ 5 animals each) revealed that the proteins encoding all genes that were enriched in the AC (2-fold or greater) were present at high levels in the AC, and most were expressed at levels higher than neighboring uterine cells (Fig. 1E, Fig. S2). TCT-1, which was highly enriched in the AC, was also at high levels in neighboring uterine cells (Fig. S3A). Notably, the DMD-3 protein (doublesex), whose encoding mRNA was extremely enriched (∼45-fold enrichment), was solely detected in the AC at the mid-L3 stage (Fig. S3B). The proteins encoded by genes whose transcripts were slightly enriched or at equivalent levels in the AC (∼1-to-2-fold enrichment versus WB) were generally present at equivalent levels to uterine cells (Fig. S2). An exception was the ZMP-1 protein (GPI anchored MMP), which was present at high levels in the AC, but was not highly enriched in the transcriptome. Lack of *zmp-1* mRNA enrichment in the AC versus WB was likely a result of high levels in the WB from *zmp-1* expression in non-uterine and non-vulval tissues (Kaletsky et al., 2018). AIR-2 (Aurora B) and PAT-3 (β integrin) proteins were also examined and both were underenriched in the AC (< 0.5 fold). Consistent with this, AIR-2 was at low-to-undetectable levels in the AC, but present in neighboring uterine cells (Fig. 1E; Fig. S3C). However, PAT-3 protein was present at high levels in the AC (Fig. S2). The low enrichment of *pat-3* was likely because of strong *pat-3* expression in body wall muscles (Gettner et al., 1995). We conclude that the AC transcriptome has a high fidelity and that AC versus WB expression identifies genes enriched in AC, but can miss genes that are also expressed highly in other tissues.

### Identification of new genes that promote AC invasion

Cell invasion requires specific gene regulatory networks, dynamic membrane protrusions, BM interactions and cell signaling (Paterson and Courtneidge, 2018; Sherwood and Plastino, 2018). To identify new genes that promote invasion, we identified AC enriched genes that encode transmembrane or secreted, cytoskeleton component/regulator, or transcription factor proteins. We filtered our search to include genes that have a mammalian ortholog and removed genes previously identified as promoting AC invasion. This led to a list of 84 genes (Table S5). We conducted an RNAi screen targeting these genes in the sensitized rrf-3 RNAi strain (Methods) (Simmer et al., 2002) and identified 13 genes whose RNAi-mediated depletion caused a significant invasion defect (Table S5).

RNAi mediated knockdown of all 13 identified genes led to modest invasion defects (∼12% to 34%, Table S5, Fig. 2A) and none caused a plate level Pvl phenotype that has been used as a selection filter for invasion defects in previous screens (Matus et al., 2010). RNAi mediated reduction of *tct-1*, which encodes the *C. elegans* ortholog of the mammalian TCTP, caused the strongest defect (∼34%). Of the remaining 12 genes, four mediate cell signaling—*mom-1* (Porcupine, Wingless signaling), *wrt-6* (Hedgehog-like ligand), *paqr-3* (Progestin and AdipoQ Receptor) and *fmi-1* (Flamingo/planar-cell polarity receptor) (Burglin and Kuwabara, 2006; Sawa and Korswagen, 2013; Svensson et al., 2011). In addition, five facilitate ER-to-plasma membrane trafficking—*dab-1* (Disabled, clathrin adaptor protein), *cyn-5* (PPIB, ER chaperon), *snb-1* (Synaptobrevin, v-Snare), and *hhat-2* (ER localized palmitoyltransferase) and *tmed-3* (TMED, Golgi organization) (Sato et al., 2014), suggesting an important role for the endomembrane system in AC invasion.

**Figure 2.**
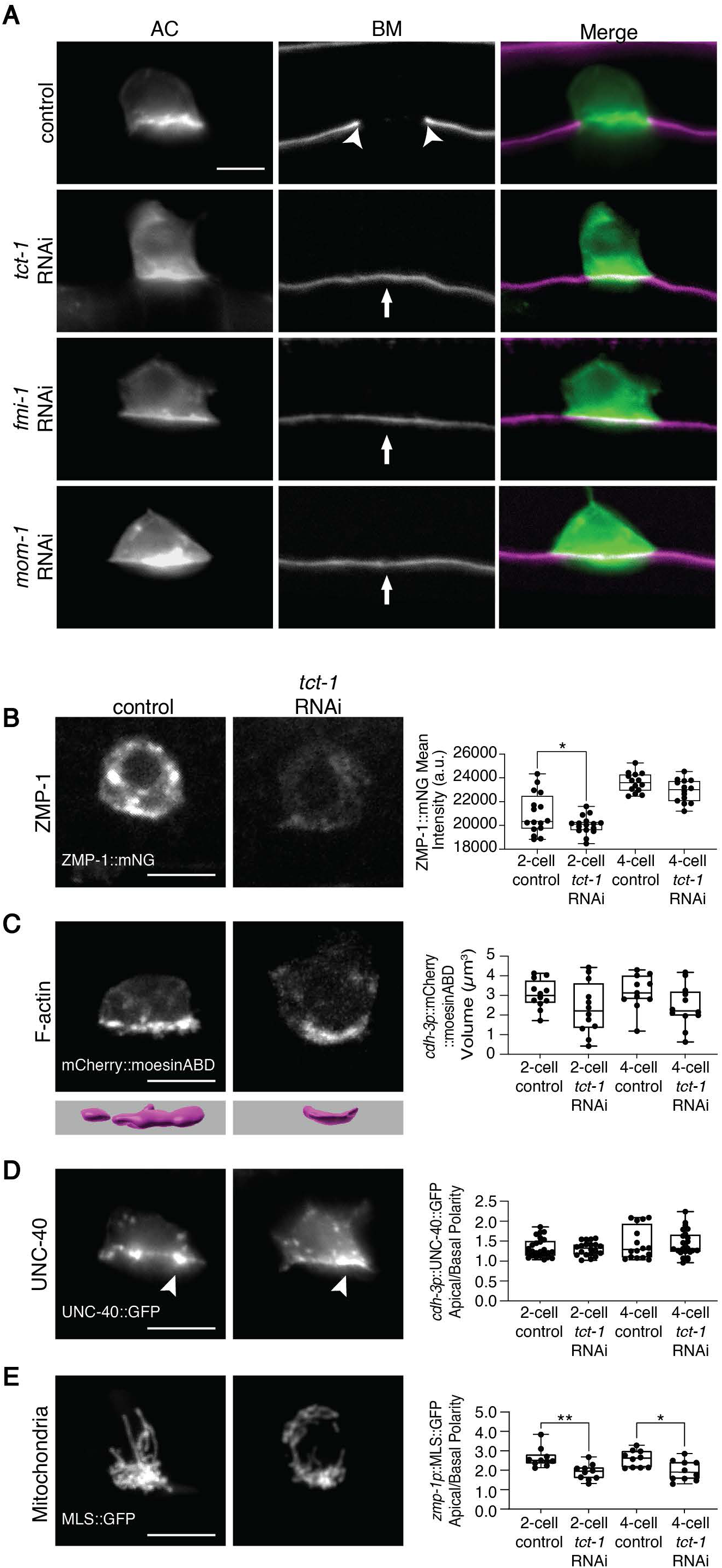
TCT-1 (TCTP) promotes AC invasion by regulating MMP expression, F-actin formation, and mitochondria localization. (A) AC (left, *cdh-3p*::GFP::CAAX) and BM (middle, *lam-1p*::LAM-1::mCherry; right, merge) at the P6.p 4-cell stage in control empty vector RNAi and RNAi targeting *tct-1*, *fmi-1*, and *mom-1*. Arrowheads mark normal BM breach and white vertical arrows show intact BM and failure to invade (B) Endogenously tagged ZMP-1::mNG levels in control and after *tct-1* RNAi treatment. Graphs show ZMP-1::mNG fluorescence mean intensity (right, boxplot n ≥ 13). (C) F-actin volume (*cdh-3p*::moesinABD::mCherry, top; isosurface renderings generated using Imaris, bottom) in control and after *tct-1* RNAi treatment. Graphs show quantification of F-actin volume (right, boxplot n ≥ 11). (D) Apical/Basal polarization of UNC-40 (*cdh-3p*::UNC-40::GFP) in control and after *tct-1* RNAi treatment. Arrowheads show enrichment of UNC-40 at AC invasive plasma membrane. Graphs show quantification of UNC-40 polarity (right, boxplot n ≥ 15). (E) AC mitochondria (*zmp-1p*::MLS::GFP) in control and after *tct-1* RNAi treatment. Graphs show quantification of mitochondrial enrichment of apical cytoplasm/basal cytoplasm (right, boxplot n ≥ 10; for graphs in B-E *p<0.05, **p<0.005, unpaired two-tailed t-tests). Quantification from these and all subsequent experiments were from two or more replicates. Scale bars: 5 µm.

TCT-1 (TCTP) encodes a multifunctional protein that interacts with many other proteins (Gao et al., 2022). TCTP has been implicated in cell invasion in human cancer cells *in vitro*, potentially through promoting a range of activities, including MMP expression and interactions with actin and microtubule regulators (Bae et al., 2015; Gao et al., 2022). To gain insight into how TCT-1 fosters invasion through BM *in vivo*, we used an AC-specific RNAi strain and confirmed that TCT-1 functions in the AC to promote invasion (Table 1). We also determined that *tct-1* RNAi knocks down AC TCT-1::mNG expression (>90% reduction, Methods). Using the *rrf-3* RNAi senstitized strain, we examined how *tct-1* loss affects several key regulators of AC invasion. RNAi mediated reduction of *tct-1* led to a decrease in the levels of the ZMP-1::mNG (MMP, Fig. 2B). There was also a modest decrease in F-actin volume at the AC’s invasive cell membrane (Fig. 2C). ZMP-1 and F-actin promote BM degradation and BM physical displacement, respectively, to remove BM during invasion (Kelley et al., 2019). Loss of *tct-1*, however, did not alter UNC-40::GFP (DCC receptor) polarization to the invasive (basal) cell membrane (Fig. 2D), but did reduce mitochondria enrichment to the invasive front (Fig. 2E). UNC-40 directs the exocytosis of lysosomes to form a large invasive protrusion at the BM breach site (Hagedorn et al., 2013; Wang et al., 2014b), while basally-localized mitochondria deliver ATP to fuel F-actin and protrusion formation (Garde et al., 2022). Together, these results suggest that TCT-1 promotes several aspects of invasion, including MMP expression, F-actin formation, and mitochondria localization, which likely accounts for the defect in AC invasion after TCT-1 loss.

**Table 1:** AC invasion after AC-specific RNAi

| Table 1: AC invasion after AC-specific RNAi |  |  |  |  |  |
| --- | --- | --- | --- | --- | --- |
|  | RNAi | Sequence ID | % ACs with incomplete invasion<sup>a</sup> | p value<sup>b</sup> | n<sup>c</sup> |
| Negative Control |  |  |  |  |  |
| 1 | L4440 (Empty Vector) | N/A | 12% | n/a | 77 |
| 2 | T444T (Empty Vector) | N/A | 5% | n/a | 20 |
| Positive Control |  |  |  |  |  |
| 3 | *fos-1* | F29G9.4 | 71% | <0.0001 | 65 |
| Large Ribosomal Proteins |  |  |  |  |  |
| 4 | *rpl-6* | R151.3 | 75% | <0.0001 | 24 |
| 5 | *rpl-4* | B0041.4 | 60% | <0.0001 | 20 |
| 6 | *rpl-38* | C06B8.8 | 60% | <0.0001 | 20 |
| 7 | *rpl-27* | C53H9.1 | 50% | 0.0002 | 20 |
| 8 | *rpl-37* | C54C6.1 | 35% | 0.01 | 20 |
| 9 | *rpl-5* | F54C9.5 | 40% | 0.03 | 10 |
| 10 | *rpl-31* | W09C5.6 | 30% | 0.04 | 20 |
| 11 | *rpl-39* | C26F1.9 | 19% | ns | 21 |
| 12 | *rpl-19* | C09D4.5 | 21% | ns | 14 |
| 13 | *rpl-33* | F10E7.7 | 16% | ns | 12 |
| 14 | *rpl-34* | C42C1.14 | 17% | ns | 12 |
| 15 | *rpl-21* | C14B9.7 | 17% | ns | 12 |
| Ribosome Biogenesis |  |  |  |  |  |
| 16 | *rbd-1* | T23F6.4 | 67% | <0.0001 | 15 |
| 17 | *W09C5.1* | W09C5.1 | 40% | 0.003 | 20 |
| 18 | *W07E6.2* | W07E6.2 | 47% | 0.004 | 15 |
| 19 | *Y48B6A.1* | Y48B6A.1 | 47% | 0.004 | 15 |
| 20 | *pro-2* | C07E3.2 | 33% | 0.05 | 15 |
| 21 | *nifk-1* | T04A8.6 | 33% | 0.05 | 15 |
| 22 | *pro-1* | R166.4 | 27% | ns | 15 |
| 23 | *W08E12.7* | W08E12.7 | 27% | ns | 15 |
| 24 | *T23D8.3* | T23D8.3 | 15% | ns | 20 |
| 25 | *fib-1* | T01C3.7 | 13% | ns | 15 |
| Endomembrane and ER Stress |  |  |  |  |  |
| 26 | *sec-61.G* | F32D8.6 | 45% | 0.002 | 20 |
| 27 | *ire-1* | C41C4.4 | 34% | 0.03 | 15 |
| 28 | *pek-1* | F46C3.1 | 20% | ns | 15 |
| 29 | *atf-6* | F45E6.2 | 7% | ns | 15 |
| Cytoskeletal and signaling |  |  |  |  |  |
| 30 | *ebp-2* | VW02B12L.3 | 40% | 0.01 | 20 |
| 31 | *tct-1* | F25H2.11 | 35% | 0.03 | 17 |
| 32 | *rap-1* | C27B7.8 | 33% | 0.05 | 15 |

<sup>a</sup> An AC specific RNAi strain was used for scoring invasion (see Methods).
<sup>b</sup> Fischer Exact 2x2 Tests were used. RNAi treatments were compared to the L4440 (empty vector) control for statistical analysis, except for *tct-1*, which was compared to the T444T empty vector control.
<sup>c</sup> Number of animals observed for each condition.

### Ribosomal proteins and ribosome biogenesis are required for AC invasion

We noted that 26 AC transcriptome enriched genes encoded translational regulators, and 12 encoded ribosomal proteins of the ribosomal large subunit (RPLs, Table S6). These observations suggest that ribosome biogenesis and protein translation may be upregulated to promote BM invasion. While ribosome biogenesis is a canonical feature of cell growth and proliferation (Donati et al., 2012), recent findings have implicated ribosome production driving EMT in cancer cell lines and in neural crest cells *in vivo* (Prakash et al., 2019). How ribosome biogenesis fuels EMT, however, is not understood.

As cells undergoing EMT require BM breaching (Kelley et al., 2014), we investigated whether ribosome biogenesis and translation are necessary for AC invasion. Using an AC-specific RNAi strain, where the AC becomes sensitive to RNAi shortly after the time of its specification (Methods), we targeted 10 AC-enriched RPLs, as well as RPL-4 and RPL-6, which are core RPLs in mouse embryonic stem cells (Shi et al., 2017). RNAi-mediated depletion of seven RPLs, including RPL-4 and RPL-6, gave strong BM invasion defects (Fig. 3A, Table 1). This suggests a general ribosome upregulation requirement, rather than a function for specific RPLs. To test the importance of ribosome production prior to invasion, we also used RNAi to deplete 10 ribosome biogenesis proteins in the AC-specific RNAi strain. These included *W08E12.7* (Pa2g4) (Tummala et al., 2016), which had high AC transcript levels (Table S1), as well as *W07E6.2* (NLE1), *Y48B6A.1* (BOP1), *pro-1* (WDR18), *pro-2* (NOC2P), and *fib-1* (FBL), which have been previously characterized as ribosome biogenesis genes in *C. elegans* (Voutev et al., 2006; Yi et al., 2015). We also targeted the *C. elegans* orthologs of the vertebrate ribosome biogenesis genes *rbd-1* (RBM19), *T23D8.3* (LTV1), *W09C5.1* (NSA2) and *nifk-1* (NIFK) (Collins et al., 2018; Kallberg et al., 2012; Pan et al., 2015; Xing et al., 2018). RNAi-mediated loss of seven ribosome biogenesis genes gave significant invasion defects (Fig. 3A, Table 1). In addition, a viable loss-of-function allele in the ribosome biogenesis gene *ddx-52* (*gc51*) (ROK1) (Itani et al., 2021) also had defects in cell invasion (Table S7). Together, these results strongly implicate ribosome production in promoting invasion after AC specification.

**Figure 3.**
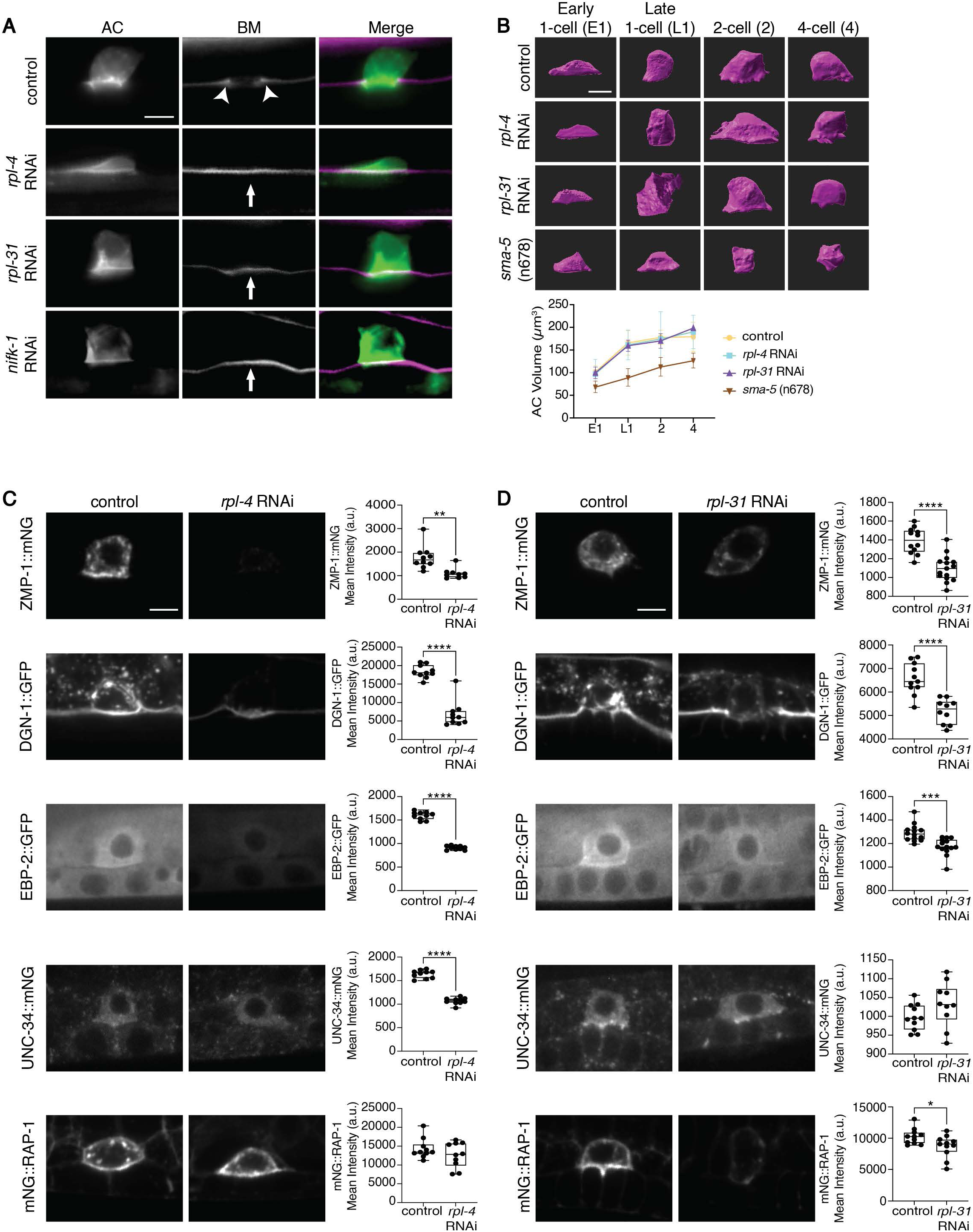
Ribosomal proteins promote invasion and the translation of pro-invasive proteins. (A) AC (left, *cdh-3p*::GFP::CAAX) and BM (middle, *lam-1p*::LAM-1::mCherry; right, merge) at the P6.p 4-cell stage in control empty vector RNAi and RNAi targeting *rpl-4*, *rpl-31*, and *nifk-1.* Arrowheads mark BM breach and white vertical arrows mark intact BM and failure to invade. (B) AC (*cdh-3p*::mCherry::PLCδ^PH^) volume from isosurface renderings of ACs from early P6.p 1-cell to 4-cell stage in control empty vector RNAi and after RNAi mediated reduction of *rpl-4* and *rpl-31*, and in *sma-*5 *(n678)* mutants. Graphs show quantification of cell volume over time (n ≥ 4 ACs per timepoint, mean ± SD, one-way ANOVA with *post hoc* Tukey’s test). (C, D) AC expression of endogenously tagged ZMP-1::mNG, DGN-1::GFP, EBP-2::GFP, UNC-34::mNG, and mNG::RAP-1 at the P6.p 2-cell stage in control empty vector RNAi and after RNAi targeting *rpl-4* (C) and *rpl-31* (D). Graphs show quantification of mean intensity (boxplot n ≥ 10 each, *p<0.05; **p<0.005; ***p<0.001, ****p<0.0001, unpaired two-tailed t-tests). Scale bars: 5 µm.

We next wanted to determine how disruption of ribosomes perturbs invasion. Loss of ribosome production can lead to nucleolar stress and p53 stabilization/activation in growing and dividing cells (Rubbi and Milner, 2003; Zhang and Lu, 2009). We thus wondered if p53 stabilization might also occur in the AC and disrupt invasion. RNAi mediated depletion of the core RPL, RPL-4, and AC-enriched RPL-37 and RPL-31, however, did not increase CEP-1::eGFP (p53 ortholog (Derry et al., 2001)) levels in the AC (Fig. S4, n=10/10 each), strongly suggesting that perturbation of AC invasion after ribosome reduction was not due to p53 activation.

We also noted that the AC size increases dramatically prior to invasion, nearly doubling in volume (Fig. 3B). As ribosome production promotes cell growth and is required for cell cycle progression and proliferation (Donati et al., 2012), we examined if ribosome biogenesis is required for AC growth and subsequent invasion. RNAi mediated loss of RPL-4 and RPL-31, however, did not reduce AC size expansion (Fig. 3B). Furthermore, we found that AC invasion proceeded normally in small body size *sma-5(n678)* animals (Watanabe et al., 2007), where the AC has a reduced size (Fig. 3B, Table S7). These results indicate that the invasion defect associated ribosome reduction is not a result of perturbations in AC growth and that AC invasion does not depend on reaching a large cell size.

Cell invasion through BM requires the translation of many new mRNA transcripts to execute invasion. To determine how reduction of ribosomes alters protein production at the time of BM breaching, we examined five endogenously tagged proteins that promote AC invasion--the Ena/VASP ortholog UNC-34, the receptor dystroglycan DGN-1, the MMP ZMP-1, the small GTPase RAP-1, and the microtubule end binding protein EBP-2 (Table 1) (Kelley et al., 2019; Naegeli et al., 2017; Sallee et al., 2018; Wang et al., 2014a). RNAi-mediated reduction of RPL-4 and RPL-31 significantly reduced all pro-invasive protein levels with the exception of RAP-1 after reduction of RPL-4 and UNC-34 after reduction of RPL-31 (Fig. 3C,D). The different effects on RAP-1 and UNC-34 might be due to the stronger reduction of RPL-4 levels after RNAi (∼50% knockdown efficiency for RPL-4 versus ∼20% for RPL-31, Methods), as the translation efficiency of distinct mRNAs can be affected by different ribosome concentrations (Mills and Green, 2017). It could also indicate distinct populations of ribosomes tailored to translate certain mRNAs (Shi et al., 2017). Overall, however, the enrichment of numerous RPLs and requirement of most for invasion, suggests a primary necessity for increased ribosome levels to support the robust translation of many pro-invasive proteins that facilitate BM invasion.

### Ribosome biogenesis occurs prior to pro-invasive protein production

We next wanted to understand how pro-invasive protein production and ribosome biogenesis are coordinated during AC invasion. We first examined the levels of nine endogenously tagged proteins within the AC that promote invasion. Levels were assessed shortly after AC specification at the early P6.p 1-cell stage until completion of invasion at the P6.p 4-cell stage (Fig. 4). The FOS-1 transcription factor, which promotes the expression of many pro-invasive genes (Medwig-Kinney et al., 2020; Sherwood et al., 2005), was present at high levels at the time of AC specification and levels were maintained throughout invasion, consistent with an early role in regulating pro-invasive gene transcription. In contrast, the other eight proteins, all of which play roles near or at the time of BM breaching, showed a significant ramp up in levels after AC specification that often increased until the time of invasion. These include the glucose transporter FGT-1, which supplies energy for BM breaching (Garde et al., 2022), as well as the integrin INA-1, the GTPase integrin regulator RAP-1, the dystroglycan receptor DGN-1, the MMP ZMP-1 (Frische et al., 2007; Hagedorn et al., 2009; Kelley et al., 2019; Naegeli et al., 2017), which regulate BM interactions, and the actin and microtubule cytoskeletal regulators Ena/VASP (UNC-34), Arp2/3 (ARX-2), and microtubule end binding protein EBP-2 (Garde et al., 2022; Kelley et al., 2019; Sallee et al., 2018; Wang et al., 2014a). Together these results indicate that there is significant pro-invasive protein production leading up to AC invasion.

**Figure 4.**
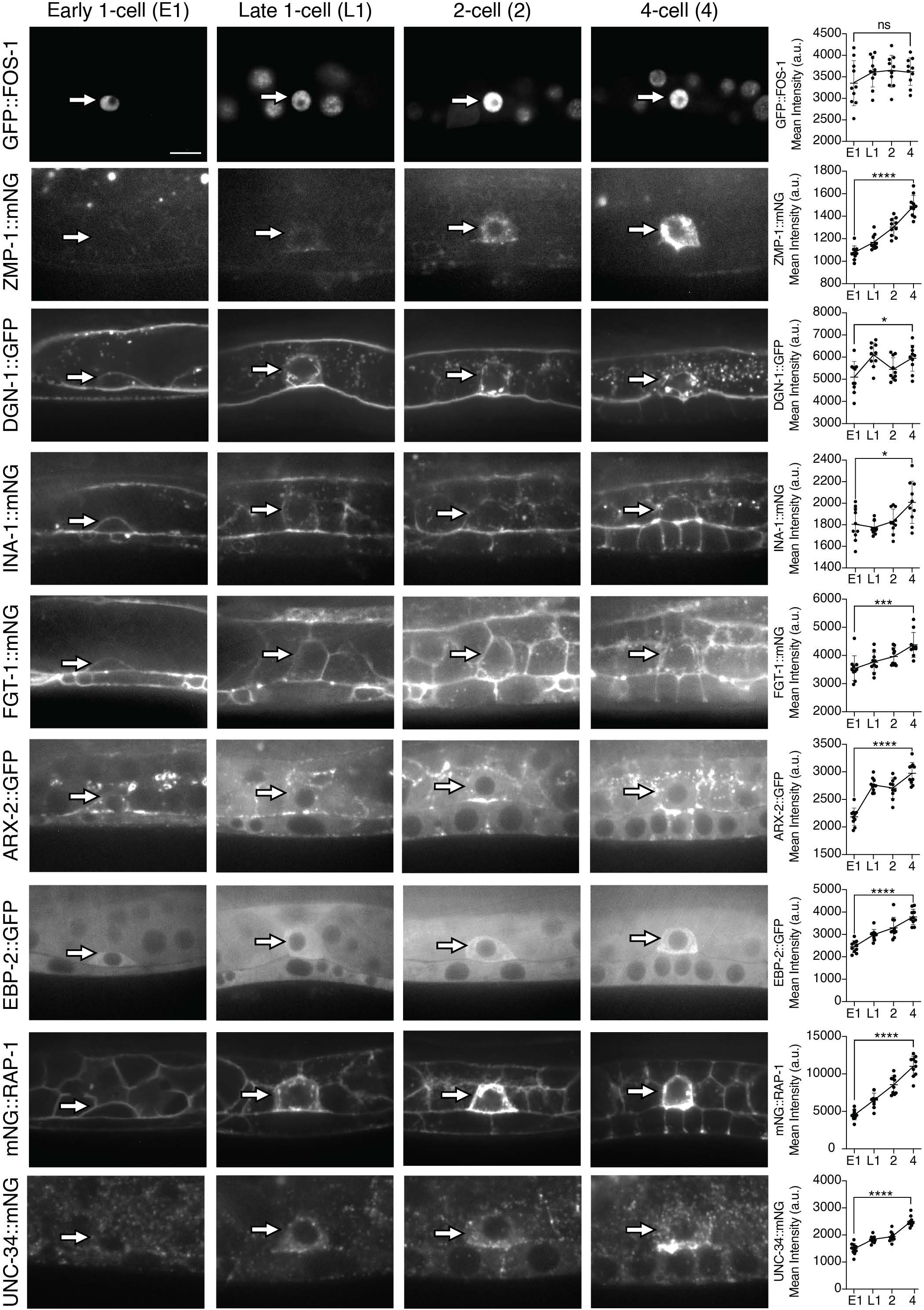
Pro-invasive proteins increase from the time of AC specification to invasion. Timeline of endogenously tagged pro-invasive protein expression in the AC (arrows) from shortly after AC specification at the early P6.p 1-cell stage (∼6 h prior to invasion) until the time when BM removal is completed at the P6.p 4-cell stage. Graphs show GFP::FOS-1 mean fluorescence protein levels did not significantly change over time, while all other pro-invasive proteins increased in levels through the time of invasion (n ≥ 10 ACs per timepoint, mean ± SD, *p<0.05; ***p<0.001, ****p<0.0001, ns, not significant, one-way ANOVA with *post hoc* Tukey’s test). Scale bar: 5 µm.

Next, we examined the timing of ribosome biogenesis and ribosome production in the AC. Nucleolar size is a strong indicator of ribosome biogenesis (Derenzini et al., 2000; Lee et al., 2012). Strikingly, the AC nucleolus showed an ∼50% increase in cross-sectional area from the early 1-cell stage (∼6 h prior to invasion) to late 1-cell stage (∼2.5 h prior to invasion) and then decreased in size during invasion (Fig. 5A). Confirming this early timing of ribosome biogenesis, levels of fibrillarin (FIB-1::eGFP), a nucleolar protein that acts as a methyltransferase for pre-rRNA processing and modification (Nguyen Van Long et al., 2022), showed the same dynamics (Fig. 5A). We also used genome editing to fuse mNG to NIFK-1, a ribosome biogenesis protein involved in pre-rRNA maturation (Pan et al., 2015), and noted a similar trend in nucleolar levels (Fig. S5A). These results suggest that there is a burst of ribosome biogenesis prior to pro-invasive protein production leading to invasion.

**Figure 5.**
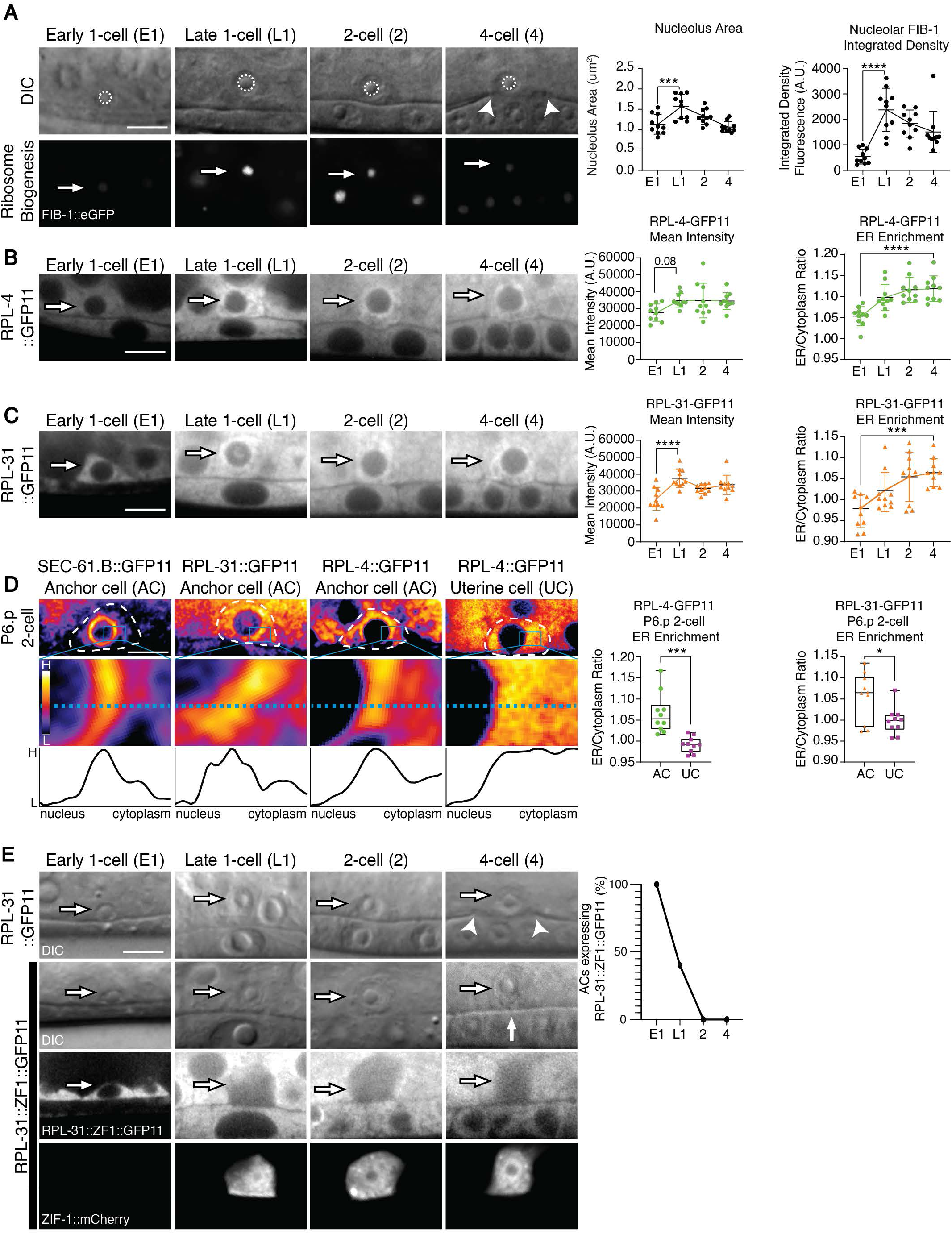
A burst of ribosome biogenesis occurs prior to invasion and ribosomes colocalize with SEC-61. (A) Timeline of nucleolus area (top row, DIC images, dotted circles) and FIB-1 levels (bottom row, arrows, *fib-1p*::FIB-1::eGFP) in the nucleolus of the AC from early P6.p 1-cell stage until time of invasion at P6.p 4-cell stage (arrowheads in top row indicate BM breach). Graphs show quantification of nucleolus area over developmental time and total fluorescence levels of FIB-1::eGFP (n ≥ 10 ACs per timepoint each, mean ± SD, ***p<0.001, ****p<0.0001, one-way ANOVA with *post hoc* Tukey’s test). (B,C) Timeline of RPL-4 (RPL-4::GFP11; *eef-1A.1p*::GFP1-10) and RPL-31 (RPL31::GFP11; *eef-1A.1p*::GFP1-10) levels in the AC (arrows). Graphs show quantification of mean fluorescence levels and ER to cytoplasm ratio (n ≥ 10 ACs per timepoint, mean ± SD, p = 0.08, ***p<0.001, ****p<0.0001, one-way ANOVA with *post hoc* Tukey’s test). (D) Spectral fluorescence-intensity map displaying the minimum and maximum pixel value range of the acquired data of SEC-61.B::GFP11, RPL-31::GFP11, RPL-4::GFP11 in the AC and in a neighboring uterine cell (UC). ACs and UC indicated with dashed white outlines. A linescan (blue dashed line, bottom panels) shows peak levels of SEC-61.B::GFP11, RPL-31::GFP11 and RPL-4::GFP11 surrounding the nucleus in the AC, whereas RPL-4::GFP11 is uniform in the cytosol of the UC. Graphs showing quantification of ER to cytoplasm ratio in P6.p 2-cell ACs and UCs expressing RPL-4::GFP11 or RPL-31::GFP11 (boxplots, n ≥ 10 each, *p<0.05; ***p<0.001, unpaired two-tailed t-test). (E) Timecourse of RPL-31::GFP11 control (DIC, top row) and RPL-31::ZF1::GFP11 animals (DIC, middle top row; fluorescence, middle bottom row). Arrows indicate the AC. All animals expressed *eef-1A.1p*::GFP1-10. RPL-31::ZF1::GFP11 was depleted upon ZIF-1 expression (bottom row, ZIF-1 visualized with *lin-29p*::ZIF-1::SL2::mCherry) and depletion caused a disruption of invasion as shown by unbreached BM (intact phase-dense line in DIC image, middle top row, white vertical arrow). In contrast, in control RPL-31::GFP11 animals the BM was cleared under the AC (phase-dense line breached, top row, arrowheads). Graph displays reduction of RPL-31::ZF::GFP11, which correlated with the beginning of ZIF-1 expression (n ≥ 10 each stage). Scale bars: 5 µm.

Visualizing ribosomes in living animals has been challenging because large fluorophores tagged to ribosomal proteins appear to interfere with the complex and tightly packed ribosome structure (Methods (Noma et al., 2017)). To overcome this problem, a split-GFP strategy has been developed to label a ribosomal protein in *C. elegans* (Noma et al., 2017). This technique uses a cell or tissue specific promoter to expresses the first ten β-strands (GFP1-10) and a ribosomal protein fused with the last β-strand (GFP11, 16 amino acids), which allows irreversible post-translational assembly of GFP to label ribosomal proteins (Cabantous et al., 2005). However, this approach has only been used with a transgene expressing the small ribosomal subunit protein RPS-18 fused to GFP11 (Noma et al., 2017), precluding clear insight into endogenous dynamics of ribosome assembly. To determine if this approach can follow endogenous ribosomal proteins, we used genome editing to tag the RPL genes *rpl-4* and *rpl-31* with GFP11 in a strain expressing GFP1-10 at high and likely saturating levels driven by the ubiquitous promoter *eef-1A.1* ((Tomioka et al., 2016), Fig S5B). Notably, RPL-4::GFP11 and RPL-31::GFP11 showed strong expression in animals and the worms were fertile and healthy (Methods). Consistent with a burst of ribosome production occurring shortly after AC specification, levels of RPL4::GFP11 and RPL-31::GFP11 increased in the AC between the early P6.p 1-cell stage and late P6.p 1-cell stage (Fig. 5B,C). Together, these results offer compelling evidence that ribosome biogenesis occurs shortly after AC specification to facilitate robust translation of pro-invasive proteins that promote BM invasion.

### Ribosomes localize to the ER prior to AC invasion and promote invasion at this time

Ribosomes are either located in the cytosol or associated with the endoplasmic reticulum (ER) where they mediate cotranslational import of transmembrane and secreted proteins through the Sec61 translocon (O’Keefe et al., 2022). We observed that both RPL4::GFP11 and RPL31::GFP11 localized strongly in a ring around the nucleus starting at the late P6.p 1-cell stage and continued this localization to the time of invasion (Fig. 5B-D, n = 10/10 each stage). This localization was specific to the AC and not seen in neighboring non-invasive uterine cells (Fig. 5D). To examine whether SEC-61 also localizes to this site, we used a split-GFP approach and endogenously tagged *C. elegans* SEC-61.B with GFP11 and crossed it into the strain expressing GFP1-10 driven by the ubiquitous promoter *eef-1A.1.* SEC-61.B::GFP11 also localized strongly to a ring around the nucleus (Fig. 5D), suggesting that many of the AC ribosomes translate proteins translocated via Sec61 into the ER. Consistent with an important role of SEC-61 and ER translocated proteins, RNAi targeting *sec-61.G* in the AC-specific RNAi strain, lead to a strong invasion defect (Table 1). To determine if ribosome translation was important for invasion at the time of ribosome localization to SEC-61/ER, we used the temporally controlled ZIF-1/ZF1 protein degradation system in which the SOCS-box adaptor protein ZIF-1 targets proteins containing the ZF1 degron motif for proteasomal degradation (Armenti et al., 2014). We tagged RPL-31 with a 36 amino acid ZF1 motif at the C-terminus followed by GFP11. We used an AC-specific promoter to express ZIF-1 fused to mCherry (*lin-29p*::ZIF-1::mCherry) and found that the ZIF-1 protein first became detectable at the late P6.p 1-cell stage when ribosomes localize to SEC-61/ER (∼2.5 hours prior to invasion, n=10/10 ACs). At this time, loss of RPL-31::ZF1::GFP11 was also first observed (n=6/10 no signal). By the P6.p 2- and 4-cell stages, RPL-31::ZF1::GFP11 was not detected in the AC (Fig. 5E, n=10/10). The late loss of RPL-31 resulted in a highly penetrant invasion defect (Table S7). Taken together these results suggest that AC ribosomes enrich at SEC-61/ER prior to invasion and that their activity is required for BM breaching at this time.

### The AC endomembrane system expands prior to invasion and is sensitive to ER stress

The localization of the ribosomes to the ER and invasion defects after SEC-61 reduction suggested an enhanced necessity for transmembrane and secreted proteins to breach the BM. Although we did not detect an enrichment of proteins with a secretion signal in our AC transcriptome, the AC produces many transmembrane and secreted proteins that mediate BM breaching (*e.g.,* integrin, dystroglycan, and MMPs, Fig. S2, Fig. 4). To examine the development of the endomembrane system after AC specification, we first examined SEC-61.B::GFP11 (translocon) and found there was a significant increase in SEC-61.B in the AC between the early and late P6.p 1-cell stage, followed by a plateauing in levels (Fig. 6A). In contrast, the ER (visualized with the ER localized, endogenously tagged protein ELO-1::mNG (Guha et al., 2020)), showed a later ramp up in levels, with the most significant ER increase occurring between the late P6.p 1-cell stage and P6.p 2-cell stage (Fig. 6B). An AC-expressed marker for the Golgi (*lin-29p*::AMAN-2(aa1-84)::mScarlet) (Sato et al., 2014), revealed an increase in Golgi stacks that mirrored ER expansion (Fig. 6C). Finally, endogenously tagged Rab-11.1 (mNG::RAB-11.1), a GTPase that marks exocytosis and recycling of vesicles at the plasma membrane (Ferro et al., 2021), was upregulated most during the time of BM breaching and enriched at the invasive front (Fig. 6D, n=10/10). Together, these results indicate a temporal expansion in the AC’s endomembrane system—from the early SEC-61 increase to import proteins into the ER, followed by the expansion of the ER and Golgi, to finally the RAB-11.1 delivery of pro-invasive proteins to the cell membrane during BM breaching.

**Figure 6.**
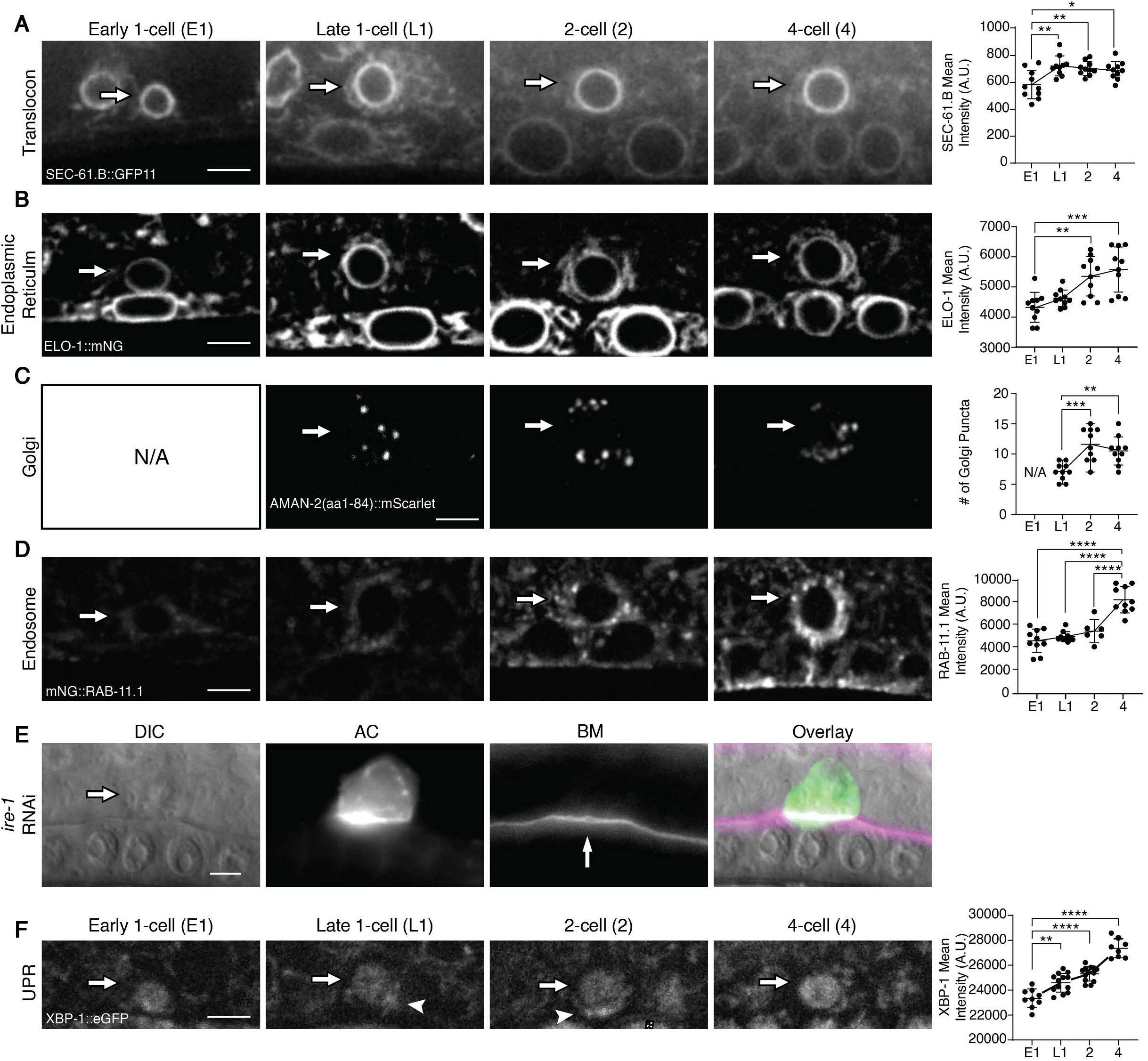
Expansion of the AC’s endomembrane system occurs prior to invasion. (A-D) A time course from the early P6.p 1-cell stage through the P6.p 4-cell stage of the AC (arrows) shows levels of the Sec61 translocon (SEC-61::GFP11; *eef-1A.1p*::GFP1-10), the ER (visualized by ELO-1::mNG), number of Golgi stacks (visualized with *lin-29*p::AMAN-2(aa1-84)::mScarlet, N/A at early P6.p 1-cell stage as *lin-29* promoter is not active at this time) and levels of recycling and secretory vesicles (mNG::RAB-11.1). Graphs show quantification of fluorescence levels and the number of golgi puncta over developmental time (n ≥ 10 ACs per timepoint, mean ± SD, *p<0.05, **p<0.005, ***p<0.001, ****p<0.0001, one-way ANOVA with *post hoc* Tukey’s test). (E) RNAi mediated reduction of the UPR sensor *ire-1* resulted in AC invasion defects as indicated by the intact BM (left, DIC, arrow AC; middle left, *cdh-3p*::mCherry::PLCδ^PH^; middle right, BM, *lam-1p*::LAM-1::GFP, white vertical arrow points to intact BM; right, overlay). (F) XBP-1::eGFP in the AC (arrows) from the early P6.p 1-cell to 4-cell stage. XBP-1::eGFP localized to the cytoplasm near site of invasion at the late 1-cell and 2-cell stages (arrowheads). Graph shows quantification of fluorescence levels over developmental time (n ≥ 10 ACs per timepoint, mean ± SD, **p<0.005; ***p<0.001; ****p<0.0001, one-way ANOVA with *post hoc* Tukey’s test). Scale bars: 5 µm

In the ER, the UPR maintains protein homeostasis by recognizing unfolded proteins and upregulating ER protein folding capacity (Read and Schroder, 2021). Cells activate the UPR during normal development and function to accommodate increased ER load or stress (Murao and Nishitoh, 2017; Wei et al., 2015). In *C. elegans* and vertebrates, three transmembrane proteins activate the UPR: the protein kinase IRE1, the protein kinase PERK, and the ER anchored transcription factor ATF6 (Wei et al., 2015). To determine if the AC requires the UPR to invade, we used an AC-specific RNAi strain to target each UPR sensor and found that loss of *ire-1* resulted in an invasion defect (Fig. 6E, Table 1). Animals harboring a deletion allele of *ire-1* (*ok799)* (Henis-Korenblit et al., 2010) also had invasion defects (Table S7). Activation of IRE-1 results in its catalyzing a non-conventional cytosolic splicing of an intron in the mRNA encoding the bZIP transcription factor XBP-1, which when translated enters the nucleus to upregulate genes that relieve ER stress (Read and Schroder, 2021). Consistent with activation of IRE-1 in the AC leading to invasion, an endogenously tagged reporter for XBP-1 (XBP-1::eGFP) localized to the nucleus of the AC, and nuclear levels increased up until the time of invasion (Fig. 6F). We also observed cytosolic localization of XBP-1 near the site of BM invasion at the late P6.p 1-cell and 2-cell stages (n=10/10 each), suggesting that ER stress is activated towards the invasive front. Taken together, these findings suggest that ribosome biogenesis and increased translation is directed in part through the AC’s endomembrane system, which expands and activates IRE1 to traffic proteins that mediate BM breaching.

## DISCUSSION

The ability of cells to invade through BM barriers plays pivotal roles in development, immune cell trafficking and metastasis. Cell invasion through BM, however, is often stochastic *in vivo* and it is difficult to faithfully recapitulate BM complexity and cellular interactions with *in vitro* invasion assays (Kelley et al., 2014; Schoumacher et al., 2013). To identify genes associated with invasive ability, expression profiling has used invasive cancer cell lines, tumor microdissection, isolating infiltrating T cells, single cell sequencing of whole tumors, and laser capture microdissection of infiltrating cancer cells (Hoelzinger et al., 2005; Kadonaga et al., 2022; Tkachev et al., 2021; Tokura et al., 2022; Zajchowski et al., 2001). While these approaches have revealed important factors that regulate cell migration through tissues and *in vitro* matrices, they have not isolated pure populations of cells invading through BM. We have taken advantage of the stereotyped timing of AC invasion in *C. elegans* to generate an AC transcriptome during BM breaching. Strongly supporting the rigor of the AC transcriptome, almost all known AC expressed regulators of invasion were present and many highly enriched, such as the transcription factors *fos-1, egl-43, nhr-67* and *hlh-2* (Medwig-Kinney et al., 2020). Furthermore, we found that *air-2* (Aurora B kinase), an essential regulator of cell division (Richie and Golden, 2005), was downregulated in the AC, which is consistent with the known requirement of G1 cell cycle arrest for AC invasion (Matus et al., 2015).

Through a focused RNAi screen of a subset of AC transcriptome enriched genes, we discovered a requirement for TCT-1 (Translationally Controlled Tumor Protein, TCTP) in AC invasion. TCTP is a multifunctional protein that is thought to regulate the stability or activity of over 100 proteins (Bommer and Telerman, 2020). TCTP’s promiscuity likely underlies its influence on numerous cellular processes, including cell growth, cell division, apoptosis, and response to cell stress (Bommer and Telerman, 2020). TCTP is overexpressed in most tumors, is associated with poor patient prognosis, and is being targeted in cancer clinical trials (Boia-Ferreira et al., 2017; Bommer, 2017; Gao et al., 2022; Karaki et al., 2017). Although strongly implicated in metastasis (Gao et al., 2022; Zhang et al., 2017), a role for TCTP in BM invasion has not been establish *in vivo*. Through endogenous tagging and AC-specific RNAi, we found that TCT-1 (TCTP) is present at high levels in the AC cytoplasm and functions within the AC to promote invasion. Loss of TCT-1 led to a reduction in F-actin within invasive protrusions, a decrease in levels of the BM degrading MMP ZMP-1, and a reduction in the enrichment of mitochondria at the invasive front (Garde et al., 2022; Kelley et al., 2019). These observations are consistent with TCTP’s multiple molecular targets and suggest that inhibiting TCTP in cancers may be effective in preventing BM invasion.

Ribosome biogenesis is an established driver of cell growth and proliferation (Donati et al., 2012). Recent studies have also linked ribosome biogenesis to EMT and the differentiation of stem cells (Prakash et al., 2019; Sanchez et al., 2016). Analysis of germ stem cell differentiation in *Drosophila* has indicated that ribosome biogenesis occurs before germ stem cell differentiation when translation levels are low and that during differentiation translation increases (Sanchez et al., 2016). As differentiation requires higher protein synthesis (Teixeira and Lehmann, 2019), it suggest that ribosome biogenesis expands translation capacity early in stem cells to allow protein translation required for differentiation (Breznak et al., 2023). We found that numerous translational regulators were enriched in the AC transcriptome, including many ribosomal large subunit (RPL) proteins. Furthermore, AC-specific RNAi knockdown of RPLs and ribosome biogenesis factors indicated that ribosome biogenesis is required to support AC invasion and that even a modest reduction in an RPL (∼20%, RPL-31) blocked invasion and reduced the translation proteins critical to BM breaching. By using live cell imaging to follow labelled ribosomes, ribosome biogenesis markers, and endogenous proteins involved with BM breaching, we discovered that a burst of ribosome biogenesis occurs shortly after AC specification and several hours prior to invasion when proteins involved with BM breaching increase in levels. Our observations suggests that early ribosome biogenesis expands the AC’s translation capacity to facilitate the subsequent translation of proteins that execute invasion. These findings may account for the role of ribosome biogenesis prior to EMT, where epithelial cells undergoing EMT, such as neural crest, breach the underlying BM to initiate their migration (Leonard and Taneyhill, 2020; Prakash et al., 2019). Interestingly, we noted differences in the effects on translation of several pro-invasive proteins after RNAi-mediated knockdown of two different RPLs. This could indicate specialized ribosomes built for the translation of different pro-invasive mRNAs (Shi et al., 2017) or that some mRNAs might have distinct translation sensitivities to ribosome concentrations (Mills and Green, 2017). Nevertheless, the disruption of AC invasion after RNAi-mediated reduction of numerous RPLs, strongly argues for a general ribosome increase to support BM invasion.

ECM proteins, proteases, and ECM receptors mediating BM breaching are translated by ribosomes associated with the ER Sec61 translocon, where they are translocated into the ER membrane or ER lumen and enter the endomembrane system (O’Keefe et al., 2022). By using a split-GFP approach to label endogenous ribosomes (Noma et al., 2017), we found that AC-ribosomes enriched at the site of Sec61/ER ∼2.5h prior to BM invasion. This timing corresponded to the ramp up in proteins trafficked by the endomembrane system, such as the MMP ZMP-1, the integrin receptor INA-1, the dystroglcan receptor DGN-1, and the glucose transporter FGT-1, and when the AC begins secreting the ECM protein hemicentin that modifies the BM prior to invasion (Morrissey et al., 2014). Correlating with this buildup of transmembrane and secreted proteins, we found that the endomembrane system expanded, initially by an increase in Sec61, followed by ER and Golgi and lastly by secretory vesicles (Ferro et al., 2021). Like B cells that differentiate into Ig-secreting plasma cells through expansion of secretory organelles (Kirk et al., 2010), these observations suggest that BM invasion requires endomembrane system enlargement to support increased production of secreted and transmembrane proteins. Notably, EMT, which requires BM breaching, is linked to increased ECM secretion as well as a sensitivity to ER stress (Feng et al., 2014). Screening of the ER stress sensors revealed a sensitivity of AC invasion to reduction in IRE1 and nuclear localized buildup of its target, the transcription factor XBP1 (Limia et al., 2019). XBP1 upregulates chaperones that promote ER protein folding and secretion and increases lipid synthesis for ER expansion (Limia et al., 2019). These observations further support the notion of an increased transmembrane and secretory load in the AC leading up to invasion. Activated IRE1 directs the splicing of an intron from XBP1 mRNA, which is then translated in the cytosol (Cox and Walter, 1996). Interestingly, cytosolic XBP1 was localized to the AC’s invasive front, suggesting that ER stress is generated from trafficking or translation near the BM invasion site. IRE1/XBP1 is strongly correlated with cancer progression and the initiation of metastasis when BM are breached in several types of carcinoma (Limia et al., 2019), suggesting a shared function for XBP1 in cell invasion through BM.

Together our observations support a model where ribosome biogenesis occurs shortly after AC specification to expand translation capacity, which allows pro-invasive protein production and concomitant expansion of the endomembrane system to deliver transmembrane and secreted proteins to the invasive front that enable BM breaching (Figure 7). We expect that the AC transcriptome will allow other genes and cell biological processes involved with BM breaching to be identified. As cell invasion has feedback and adaptive mechanisms that ensure robustness (Parlani et al., 2022), focused screens in mutant backgrounds might be an especially powerful approach to reveal mechanisms underlying invasion and new strategies for targeting invasive activity in cancer.

**Figure 7.**
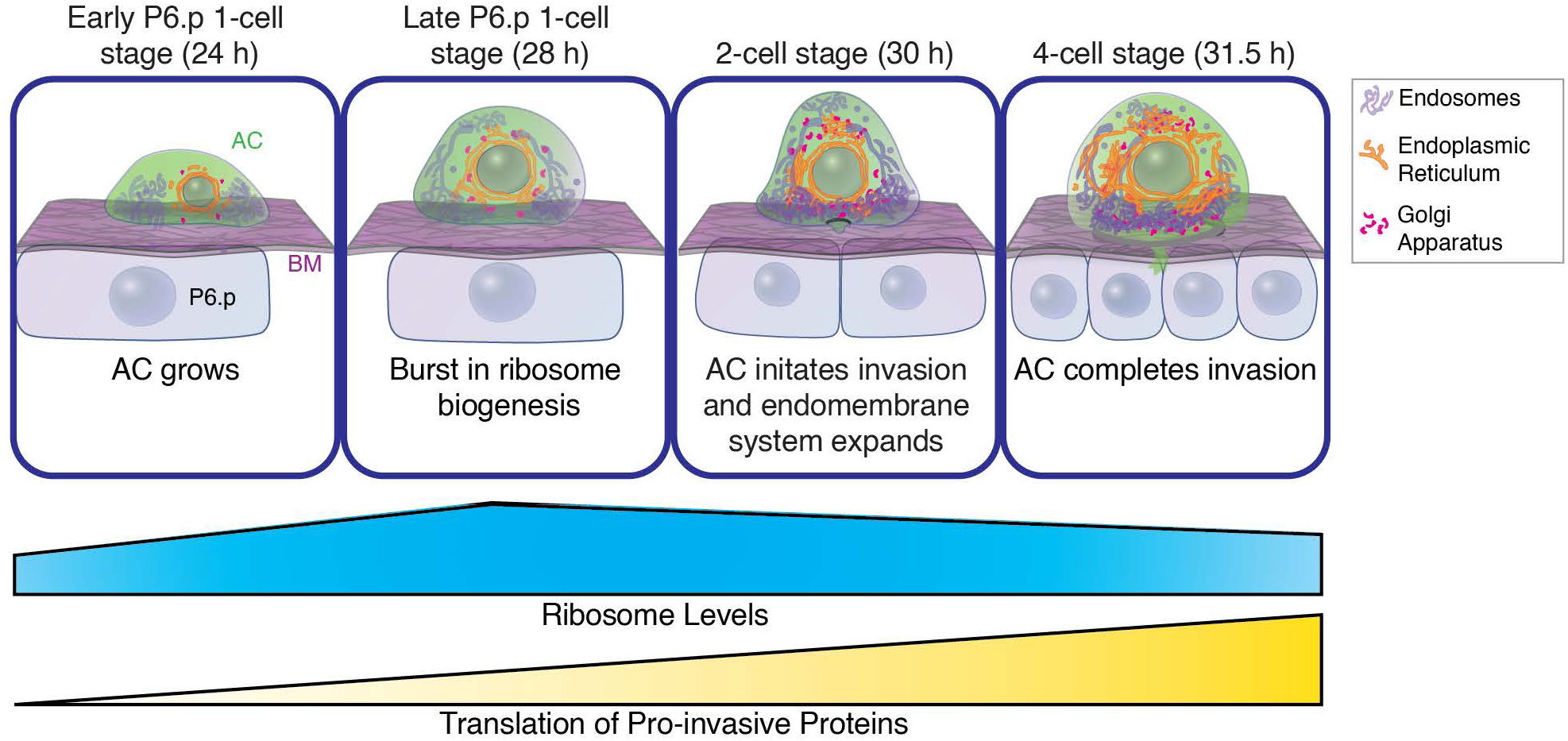
Ribosome biogenesis precedes expansion of the endomembrane system and drives pro-invasive protein translation. Schematic diagram of the AC from the time of AC specification at the early P6.p 1-cell stage to completion of AC invasion at the P6.p 4-cell stage. The timing of AC invasion are staged by the divisions of the vulval P6.p cell. Ribosome levels increase dramatically after the AC is specified and peak at the late P6.p 1-cell stage. The endomembrane system expands after the burst of ribosome biogenesis. Increased ribosome levels support pro-invasive protein translation essential for BM invasion, including proteins trafficked through the endomembrane system. Timeline in h post-hatching at 20°C shown.

## ACKNOWLEDGEMENTS

We thank the Duke University SOM Sequencing and Genomic Technologies Shared Resource, which provided DNA sequencing and flow cytometry and M. Crowder for *ddx-52(gc51)*. Some strains were provided by the Caenorhabditis Genetics Center, which is funded by National Institutes of Health Office of Research Infrastructure Programs [P40 OD010440].

## AUTHOR CONTRIBUTIONS

Conceptualization: D.S.C. and D.R.S.; Methodology: D.S.C., I.W.K., Q.C., L.C.K. and D.R.S.; Validation: D.S.C. and I.W.K.; Formal analysis: D.S.C., I.W.K.; Investigation: D.S.C., I.W.K., Q.C., K.P., and D.R.S.; Resources: D.R.S., A.G., D.Q.M., S.P., S.Y., T.V.G., A.M.P., B.G.; Data curation: D.S.C and I.W.K.; Writing - original draft: D.S.C., D.R.S.; Writing-review & editing: D.S.C., I.W.K., and D.R.S.; Visualization: D.S.C. and I.W.K. Supervision: D.R.S.; Project administration: D.R.S.; Funding acquisition: D.R.S.

## MATERIALS AND METHODS

### *C. elegans* strains and maintenance

*Caenorhabditis elegans* were grown under standard conditions at 18°C or 20°C on nematode growth media (NGM) and fed *Escherichia coli* OP50. N2 Bristol strains were used as wild-type (Brenner, 1974). All the animals used in this study were hermaphrodites and were scored at the time of anchor cell (AC) invasion during the L3 stage as described previously (Sherwood et al., 2005). For the purpose of RNAi experiments and some developmental time courses, L1 synchronization was performed with hypochlorite treatment (Stiernagle, 2006). In texts and figures, promoter-driven transgenes are denoted with a “p” following the gene’s name, and endogenously tagged proteins are denoted with a “::” annotation. The following alleles and transgenes were used in this study: *air-2(ie31[degron::GFP::air-2])* I, *bmd15[eef-1A.1p::GFP1-10::unc-54 3’ UTR; myo-2p::mCherry::3xHA::tbb-2 3’ UTR]* I, *hlh-2(bmd90[LoxP::GFP::HLH-2])* I, *rab-11.1(qy190[mNG::rab-11.1])* I, *rpl-4(qy128[rpl-4::GFP11])* I, *rpl-31(qy110[rpl-31::GFP11])* I, *rpl-31(qy189[rpl-31::ZF1::GFP11]* I, *tct-1(qy161[tct-1::mNG])* I, *qy121[eef-1A.1p::GFP]* I, *ddx-52(gc51)* I, *ebp-2(wow47[ebp-2::GFP::3xFLAG])* II, *egl-43(bmd88[LoxP::GFP::EGL-43])* II, *ire-1(ok799)* II, *ptp-3(qy47[ptp-3::mNG])* II, *qyIs17[zmp-1p::mCherry]* II, *rrf-3(pk1426)* II, *fgt-1(qy65 [fgt-1::mNG*]) II, *qyIs23[cdh3p::mCherry::PLC*δ*^PH^]* II, *unc-119(ed4)* III, *unc-119(tm4063)* III, *ina-1(qy23[ina-1::mNG])* III, *pat-3(qy36[pat-3::mNG])* III, *nifk-1(qy126[nifk-1::mNG*]) III, *sma-4(e729)* III, *ten-1(qy56[ten-1::mNG])* III, *wgIs506[xbp-1::TY::eGFP::3xFLAG + unc-119(+)]* III, *zif-1(gk117)* III, *zmp-1(qy17[zmp-1::mNG])* III, *eif-1.A(qy90[eif-1.A::mNG])* IV, *elo-1(qy97[elo-1::mNG])* IV, *him-8(e1489)* IV, *lin-3(cp226[lin-3::mNG-C1::3xFLAG])* IV, *nhr-67(syb509[nhr-67::GFP])* IV, *qyIs10[lam-1p::lam-1::GFP]* IV, *rap-1(cp151[mNG-C1::3xFLAG::rap-1])* IV, *sec-61.B(shy61[sec-61.B::GFP11x2])* IV, *arx-2(cas607[arx-2::GFP])* V, *fos-1(bmd138[LoxP::GFP::FOS-1])* V, *lag-2(cp193[lag-2::mNG::3xFLAG])* V, *ot932[dmd-3::GFP::3xFLAG]* V, *qyIs49[guk-1p::guk-1::YFP]* V, *qyIs127[lam-1p::lam-1::mCherry + unc-119(+)]* V, *rde-1(ne219)* V, *snb-1(qy164[snb-1::mNG])* V, *unc-34(ljf3[unc-34::mNG[C1]^3xFlag::AID]*) V, *dgn-1(qy206[dgn-1::GFP])* X, *qyIs166[cdh-3p::GFP::CAAX + unc-119(+)]* X, *sdn-1(qy29[sdn-1::mNG])* X, *sma-5(n678)* X, *knuSi221[fib-1p::fib-1(genomic)::eGFP::fib-1 3’ UTR + unc-119(+)]*, *qyEx603[lin-29p::AMAN-2(aa1-84)::mScarlet]*, *qyIs66[cdh-3p::unc-40::GFP], qyIs24[cdh-3p::mCherry::PLC*δ*^PH^], qyIs287[myo-3p::mCherry]*, *qyIs362[lin-29p::GFP]*, *qyIs463[lin-29p::ZIF-1::SL2::mCherry]*, *qyIs550[zmp-1p::MLS::GFP], qyIs57[cdh-3p::mCherry::moesinABD + unc-119(+)], qyIs102[fos-1ap::rde-1(genomic) + myo-2::YFP + unc-119(+)], wgIs793 [cep-1::TY1::eGFP::3xFLAG + unc-119(+)]*. See Table S8 for details of strains generated and used in this study.

### FACS isolation of ACs

ACs expressing *lin-29p::GFP* were dissociated using a method similar to those described previously (Zhang and Kuhn, 2013). Briefly, a synchronized population of worms were plated on 100mM NEP plates seeded with NA22 bacteria and collected once they reach the P6.p 2-cell stage (mid L3 larval stage). An SDS-DTT (20mM HEPES pH 8.0, 0.25% SDS, 200mM DTT, 3% sucrose) solution was used to begin breaking down the cuticle, followed by 15mg/mL Pronase E treatment and mechanical disruption to further degrade the cuticle and release cells. Cells were settled in ice-cold L-15/Fetal Bovine Serum (FBS) solution and then passed through a 5 μm filter to remove any large debris. Cells were resuspended in ice-cold PBS + 2% FBS + 10 mM EDTA and sorted using BD FACSDiva equipped with a 488 nm argon laser. Emission filters for GFP were 530 ± 30 nm. Egg buffer was used to flush the machine prior to sorting. Gates were set using N2 cell suspension samples. 15,000-150,000 fluorescently labeled GFP+ events were sorted each session. N2 whole body control cells were collected in the same way as the AC but were not sorted using FACS.

### RNA isolation, amplification, library preparation, and sequencing

RNA was collected and prepared from FACS isolated GFP+ cells and wild-type (N2) cells. GFP+ cells collected from separate FACS sessions were combined so that each library contained 80,000-150,000 ACs per library. RNA was extracted using a modified protocol based on a phenol-chloroform extraction method (Roy et al., 2020). Agilent 2100 Bioanalyzer System was used to assess the quality and quantity of the RNA. We considered the RNA to be of good quality if its RNA integrity number was at least 7. cDNA was synthesized from 8-10ng of good quality RNA using the Ovation RNA-Seq Sytem V2 Kit (Part No. 7102) according to the manufacturer’s suggested practices. The generated cDNA was sheared to ∼300 bp using Covaris E220 Focused-Ultrasonicator. Libraries were prepared using KAPA hyper prep kit. Libraries were sequenced using Illumina HiSeq 4000. We obtained 40-57 million (47,686,916 average) reads for each sample and mapped them to the *C. elegans* genome (WBcel235) (http://ftp.ensembl.org/pub/release-107/fasta/caenorhabditis_elegans/dna/).

### RNA-seq data analysis

FASTQC was used to check the quality of the raw sequence data (Andrew, 2010). Trim Galore was used to remove any low-quality bases and adapter sequences (Krueger et al., 2021). The trimmed reads were mapped to the *Caenorhabditis elegans* genome (WBcel235) using STAR alignment (Dobin et al., 2013). The genomic alignment was run using the following parameters: outFilterMultimaoNmax 1, outSAMstrandField intronMOtif, alignSJoverhangMin 500. RsEQC was used to check for uniform coverage of reads (Wang et al., 2012). HTSeq was used to quantify gene expression data (Putri et al., 2022). Downstream Differential Expression analysis was performed in R version 4.0.2 (R_Core_Team, 2021) using DESeq2 v.1.30.1 (Love et al., 2014). Genes that did not contain at least 10 reads in a single AC library were filtered out. We identified 1502 AC expressed genes with a log2fold change greater than or equal to 1.0 compared to the WB libraries and a Benjamini–Hochberg adjusted p-value less than or equal to 0.1 to be significantly elevated within the AC compared to the WB (Table S2). See Table S1 for complete AC and whole-body transcriptomes.

### Gene set enrichment analysis

For gene set enrichment analysis, we selected genes with a Benjamini–Hochberg adjusted p-value less than or equal to 0.1 and a log2fold change greater than or equal to 1.5, resulting in 1189 genes. These enriched genes were then analyzed using Database for Annotation, Visualization, and Integrated Discovery (DAVID), which compared all of the genes identified in our WB transcriptome to the 1189 genes to determine important biological pathways and significant genes with the Functional Annotation Table tool (Dennis et al., 2003). WormBase ParaSite Biomart (Howe 2017) was used to identify genes with a human ortholog and identify genes that are included in the following Gene Ontology terms: protein targeting to membrane (GO:0006612), cytoskeleton (GO:0005856), DNA-binding transcription factor activity (GO:0003700), intracellular membrane-bound organelle (GO:0043231) (Howe et al., 2016; Howe et al., 2017).

### Construction of genome edited strains

CRISPR-Cas9 mediated genome editing with a self-excising hygromycin selection cassette (SEC) (Dickinson et al., 2015; Keeley et al., 2020; Meca-Cortes et al., 2017) was used to insert mNG in frame into the open reading frame of genes to endogenously tag the encoded proteins and visualize their levels under native regulatory control. Briefly, *tct-1*, *rab-11.1*, *snb-1*, *eif-1.A*, *nifk-1*, *elo-1,* and *lin-3* were tagged at their C-termini, with a double linker fused to mNG, which was inserted directly in front of the stop codon. To visualize endogenous DGN-1 levels, *dgn-1* was tagged at the C-terminus with a double linker fused to GFP inserted directly in front of the stop codon. To generate the *rpl-4* and *rpl-31* split-GFP constructs, a linker fused to GFP11 was inserted at the C-terminus directly in front of the stop codon. Endogenous *sec-61.B* was tagged with GFP11x2 following previously described protocols (Ghanta et al., 2021). For all constructs, 2-3 KB of DNA centered around the insertion site was amplified from N2 genomic DNA and cloned into an intermediate vector (TOPO) to be used as a template for homology arms. Primers were used to introduce silent point mutations adjacent to the Cas9 cut site. The mutated homology arms were inserted into the repair plasmids mNG-SEC-LL (Keeley et al., 2020), GFP-SEC (Dickinson et al., 2013), or GFP-11-SEC (this study) using New England Biolabs Gibson assembly and confirmed with PCR. One or two short guide RNA (sgRNA) plasmids were generated for each target by cutting plasmid pDD122 with Nhel and EcoRV, then using New England Biolabs Gibson assembly to insert the respective sgRNA sequences into the plasmid. The *lin-3* guide sequence was inserted into pDD162 (*eef-1A.1p::*Cas9 + empty sgRNA, Addgene plasmid #47549) using Q5 site-directed mutagenesis (New England Biolabs). The sgRNA targeting sequences are listed in Table S9.

To generate the *eef-1.a.1p*::GFP construct, Gibson Assembly was used to insert the *eef-1a.1* promoter and GFP into the Multiple Cloning Site vector pCFJ352. CRISPR-Cas9 mediated recombination was used to insert *eef-1a.1p::GFP* into the standard MosSCI insertion site ttTi4348 on Chromosome I (Dickinson and Goldstein, 2016). Oligonucleotide sequences are listed in Table S9.

Microinjection of the constructs, hygromycin selection, and excision of the SEC cassette were carried out as previously described (Dickinson and Goldstein, 2016; Keeley et al., 2020). Split-GFP constructs for *rpl-4* and *rpl-31* and guides were injected into animals with ubiquitous somatic expression of GFP1-10 (*eef-1a.1::GFP1-10)* and a pharyngeal promoter driving mCherry (*myo-2p::mCherry)*. Since GFP1-10 is not fluorescent until assembled with GFP11, the red pharyngeal marker was used to track GFP1-10 positive animals. The *eef-1A.1p*::GFP construct was used to confirm that the *eef-1A.1* promoter expresses GFP at similar levels in the AC from the time of specification through the time of BM breaching.

The *sec-61.B* repair template with GFP11x7 and Cas9 protein (IDT) duplexed with tracrRNA, and targeting crRNA (TCGTGGGCTGCGTCCAGAAT, IDT) were injected directly into the gonad of MTS606, a strain carrying an integrated array of *mig-13*p::GFP1-10 (*shyIs32*). For the repair template, 35bp homology arms flanking the GFP11x7 were PCR amplified and gel purified to make the final concentration of 25ng/µL in the injection mixture. The mNG and GFP repair constructs and guides were injected into a wild-type background (N2). To verify the proper insertion of the constructs, homozygous animals with fluorescence were confirmed with PCR genotyping, and the site of the insertion was sequenced with the exception of UNC-34 (see Table S9 for a list of genotyping primers). Endogenous *sec-61.B* was confirmed using Sanger sequencing with primers flanking the KI site. mNG::RAB-11.1 worms were maintained as heterozygotes because homozygous worms were sterile. Attempts to tag *rpl-4, rpl-6, rpl-31,* and *rpl-37* directly with mNG directly were not successful as lines were not viable.

### RNA interference and AC specific RNAi sensitive strain

RNAi clones originated from either an RNAi library constructed by the Ahringer Lab (Kamath et al., 2003), the *C. elegans* ORF-RNAi V1.1 library (Open BioScience) (Rual et al., 2004), or were generated in this study (Table S9). RNAi clones were sequenced using M13 Forward primer and sequences were aligned to the *C.elegans* genome using BLAST to confirm the correct clone. Correctly sequenced clones were frozen and used in subsequent experiments. RNAi was generated for *pro-1* in the L4440 vector using New England Biolabs Gibson assembly. For all RNAi screening experiments in the L4440 vector, an empty L4440 RNAi vector was used as a negative control and the RNAi clone that targets the transcription factor *fos-1*, which is causes AC invasion defects after RNAi-mediated knockdown (Sherwood et al., 2005), was used as a positive control. RNAi was generated in the more efficient T444T RNAi vector for *tct-1* (Sturm et al., 2018) using New England Biolabs Gibson assembly. The empty vector T444T was used as a negative control. Both *pro-1* and *tct-1* RNAi plasmids were transformed into *E. coli* HT115(DE3) bacteria. See Table S9 for a list of primers used to generate RNAi clones in this study.

RNAi was delivered to animals by feeding *E. coli* strain HT115 (DE3), which expresses double-stranded RNA (Timmons et al., 2000). RNAi cultures were grown for 12-16 h at 37 °C in lysogeny broth with 1 μl/ml ampicillin. To induce transcription of RNAi vector expression, cultures were grown for an additional hour after the addition of 1 mM Isopropyl b-D-1-thiogalactopyranoside (IPTG). RNAi cultures were seeded onto 60 mm NGM plates with topically applied 1 mM IPTG and 100 mg/ml ampicillin and left at room temperature overnight to allow for further induction. A synchronized population of L1 Larvae (Porta-de-la-Riva et al., 2012) was plated on the RNAi feeding plates and grown at 20°C for 36-40 h before AC invasion was scored at the L3 stage.

Sensitized RNAi screening using the *rrf-3 (pk1426)* mutant background was performed as previously described (Matus et al., 2010). AC-specific RNAi experiments were preformed as previously described using *rrf-3 (pk1426); qyIs102[fos-1p::rde-1; myo-2::GFP]; qyIs10[lam-1p::lam-1:GFP]; rde-1(ne219)* (Hagedorn et al., 2009). This strain uses animals possessing a null mutation of *rde-1*, which is required for RNAi sensitivity (Tabara et al., 1999). A functional copy of RDE-1 is expressed in the AC at high levels with the *fos-1* promoter beginning at the time of AC specification (∼6 h prior to invasion (Medwig-Kinney et al., 2020)) and at low levels in neighboring uterine cells (the uterine cells do not influence AC invasion (Sherwood and Sternberg, 2003)), thus restoring RNAi in the AC to determine site of action of the gene and avoid earlier requirements of the gene in development.

### Assessment of AC invasion

Anchor cell invasion was scored as previously described (Hagedorn et al., 2009; Sherwood and Sternberg, 2003). Briefly, animals were scored for invasion at the VPC P6.p 4-cell stage when BM invasion is completed in 100% of wild-type animals. Wild-type invasion was defined as a breach in the BM that is the width of the ACs basal surface. Invasion defects are defined as a fully intact BM (blocked invasion) or a smaller than normal breach in the BM that is less than the width of the ACs nucleus (partial invasion). The BM was either assessed examining the presence or absence of the phase-dense line under the AC by DIC microscopy or by examining animals expressing GFP or mCherry tagged laminin (*lam-1p::lam-1::GFP* or *lam-1p::lam-1::mCherry*).

### Microscopy and image acquisition

Confocal Microscopy images were acquired using a Zeiss AxioImager microscope controlled by Micromanager software (Edelstein et al., 2010). The Zeiss AxioImager was equipped with a Yokogawa CSU-10 spinning disc confocal, a Zeiss 100x Plan-Apochromat 1.4NA oil immersion objective or a Zeiss 40× Plan-Apochromat 1.4-NA oil immersion objective and a Hamamatsu Orca-Fusion sCMOS camera or ImageEM EMCCD camera. Micromanager imaging software was used for microscopy automation. Animals were mounted into a drop of M9 onto 5% agar pads containing 0.01M sodium azide and a coverslip was placed on top. AC invasion staging was completed using 100x objective (1000X magnification) to identify the developmental stage of the P6.p cell or its descendants in the mid L3 larva. Images of dissociated cells (Fig. 1C), previously identified AC regulators present in AC transcriptomes (Fig. S2), FIB-1::eGFP fluorescence levels (Fig. 5A), and DIC images of nucleolar size (Fig. 5A), were acquired on a Zeiss AxioImager A1 microscope with a Zeiss 40x Plan-APOCHROMAT objective or a Zeiss 100x Plan-APOCHROMAT objective and Zeiss AxioCam 305 mono CMOS camera controlled by Zeiss ZEN microscopy software (Zeiss Microimaging). Three-dimensional reconstructions were built from confocal z-stacks using Imaris 9.9 (Bitplane).

### GFP11 CRISPR Strain Viability Analysis

We assessed the phenotypic normalcy of genome edited GFP11 ribosomal strains by confirming the absence of any plate-level phenotypes (e.g., uncoordinated (Unc), protruding vulva (Pvl), dumpy (Dpy), and sterile (Ste)). To assess viability, five genome edited L4 larval stage animals and five N2 L4 animals were plated on *E. coli* OP50 containing 60mm agar plates and allowed to grow until starvation. One day prior to starvation, five more L4 animals were picked to new plates for a second generation. This was repeated for a third generation. If no differences were seen in growth, the worms were considered to have N2 viability.

### Analysis of F-actin Volume

F-actin volume was determined as described previously (Kelley et al., 2019). Briefly, a confocal z-stack (0.37 µm step size, 20-28 slices) was taken through ACs expressing *cdh-3p::mCherry::moesinABD*. Imaris 9.9 was used to generate 3D reconstructions of the F-actin using the Isosurface rendering module using an absolute threshold surrounding the F-actin at the invasive surface with a 0.124 µm surface detail. Total volumetric measurements were then made.

### Determination of Polarization of UNC-40 and mitochondria in the AC

To determine polarization of UNC-40 within the AC, a 6-slice sum projection of a confocal z-stack (0.37 µm step size) through the central region of the AC expressing *cdh-3p::UNC-40::GFP* was collected. Polarization within the AC was measured as the ratio of the mean fluorescence intensity in the invasive (basal) versus the AC’s noninvasive (apical and lateral) plasma membranes using a five-pixel-wide line manually drawn in Fiji/ImageJ. Mitochondrial polarization was determined by building a 28-slice sum projection from a confocal z-stack (0.37 µm step size) through the entire AC (with background subtraction, 20 pixel rolling ball radius) expressing *zmp-1p::MLS::GFP*. The AC was separated by hand into two equal basal and apical regions by drawing ROIs. The mean fluorescence intensity was measured, and the mitochondrial enrichment was determined as the ratio of the mean fluorescence intensity of the invasive (basal) versus the AC’s noninvasive (apical) signal.

### Quantification of AC Size

Using AC expressing plasma membrane localized mCherry (*cdh-3p::mCherry:: PLC*δ*^PH^*) a confocal z-stack was collected through the entire AC (14-24 total slices depending on size of the AC, 0.37 µm step size). 3D isosurface renderings of the AC were generated in Imaris 9.9 using an absolute threshold and 0.124 µm surface detail. The total volumetric measurements were made for N2 worms, *rpl-4* RNAi treated worms, *rpl-31* RNAi treated worms, and *sma-5(n678)* mutant worms. *Sma-5(n678)* mutants crossed with worms expressing AC-specific mCherry (*cdh-3p::mCherry:: PLC*δ*^PH^*) were confirmed to be *sma-5* mutants by PCR genotyping (see Table S9 for oligonucleotides used).

### Scoring of AC CEP-1::eGFP (p53)

L1 synchronized CEP-1::eGFP animals were plated on empty vector control or RNAi targeting *rpl-4, rpl-31,* or *rpl-*37, then imaged at the P6.p 2-cell stage for the presence or absence of CEP-1::eGFP signal in the AC.

### Analysis of Invasive Protein Levels in the AC

To measure fluorescent protein levels in the AC, the mean fluorescence intensity was determined to control for variability in cell size. For the analysis of endogenous levels of pro-invasive proteins over time (Fig. 4) a 6-slice sum projection of a confocal z-stack (0.37 µm step size) through the central region of the AC was collected for each time point. All developmental time points for each strain were acquired with the same laser power and exposure time to allow for direct comparison. Mean fluorescence intensity was calculated from sum projections by outlining the AC manually as the ROI. The same imaging and analysis procedure was used in the experiments targeting ribosomal protein genes and *tct-1* with RNAi to determine the effects on endogenously tagged pro-invasive protein levels (Fig. 2B and Fig. 3C,D).

### Ribosome Biogenesis markers, RPL-4 and RPL-31 levels, and ER Enrichment

To examine ribosome biogenesis in the AC, a single confocal slice through the center of the AC was taken of ribosome biogenesis markers *fib-1p::*FIB-1::eGFP and NIFK-1::mNG. Levels were then determined by setting a threshold that outlined the intense signal in the nucleolus of the AC where the integrated density of fluorescence was then calculated. For nucleolus area measurements, DIC images the nucleoli of N2 worms were manually outlined and area was measured in FIJI. Levels of reconstituted split-GFP RPL-31::GFP11 and RPL-4::GFP11 were determined by taking a single confocal slice through the center of the AC and drawing a region of interest around the AC and measuring the mean fluorescence of the total signal. RPL-31::GFP11, and RPL-4::GFP11 enrichment at the site of SEC-61 around the nucleus was assessed by measuring the mean fluorescence intensity of a five-pixel-wide line drawn within the intense signal around the nucleus (site of strongest Sec61 localization) divided by the mean fluorescence intensity of a five-pixel-wide line within the cytoplasm just under the plasma membrane. Line-scan graphs showing RPL-4::GFP11, RPL-31::GFP11, SEC-61.B::GFP11 were generated by drawing a 2.5 µm long line within each respective representative image. The intensities of the pixel values were measured using the “Plot Profile” function in FIJI

### RPL-31::ZF::GFP11 Degredation

AC-specific ZF1 mediated protein degradation experiments were completed in a *zif-1*(*gk117*) null mutant background to avoid degrading RPL-31 in the embryo when endogenous ZIF-1 protein is normally expressed (Armenti et al., 2014). We rescued ZIF-1 expression only in the AC using the *lin-29p*::ZIF-1::SL2::mCherry, which initiates expression just before the late P6.p 1-cell stage (about 2.5 hours prior to invasion).

### Endomembrane Expansion and UPR Activation

For analysis of SEC-61.B::GFP11 and ELO-1::mNG, fluorescence levels in the AC, a 5-slice sum projection of a confocal z-stack (0.37 µm step size) was generated through the central region of the AC for each time point. For mNG::RAB-11.1, a single confocal slice was used. For each, a region of interest was manually drawn around the AC, and the total signal was measured. To count Golgi puncta, a sum projection of of *lin-29p::AMAN-2(aa1-84)::mScarlet* expression was generated from confocal z-stacks (0.37 µm step size, 20-28 slices) that sectioned through the entire AC. The number of Golgi puncta was determined using the “Spots” module in Imaris 9.9. The following parameters were used to count the puncta present: XY diameter was set to 0.5µm, Filter type “quality”, and boundary was set to 75. A single confocal slice was used to measure the mean intensity of XBP-1::eGFP fluorescence in the AC. A region of interest was manually drawn around the AC, and the total signal was measured to determine mean intensity.

### RNAi Knockdown Efficiency

To assess the knockdown efficiency of *tct-1* and *nifk-1* RNAi, animals expressing endogenous TCT-1::mNG or NIFK-1::mNG were plated onto empty vector RNAi bacteria and *tct-1* and *nifk-1* RNAi as L1 larva and allowed to grow at 20°C until they reached P6.p 2-cell stage or P6.p 4-cell stage. Mean fluorescence intensity of TCT-1::mNG was quantified from a single confocal slice using Fiji to manually draw a region of interest around the AC. RNAi targeting *tct-1* resulted in a 90% knockdown of TCT-1::mNG signal (n ≥ 7 P6.p 2-cell ACs). Mean fluorescence intensity of NIFK-1::mNG was measured from a single slice by setting a threshold that outlined the signal in the nucleolus of the AC and RNAi treatment against *nifk-1* resulted in a 91% knockdown (n ≥ 10 P6.p 2-cell ACs). RNAi knockdown of all RPL’s and ribosome biogenesis factors caused delayed growth and some L2 developmental arrest as previously described after protein synthesis inhibition (Dalton and Curran, 2018). We thus only scored animals escaping L2 arrest that developed to the L3 stage. To optimize *rpl-4* and *rpl-31* RNAi knockdown in whole body treatement RNAi, we performed timed RNAi plating to reduce or avoid L2 arrest. Split-GFP RPL-4::GFP11 and RPL-31::GFP11 worms were synchronized as L1 larvae, grown on empty vector RNAi, and then placed on *rpl-4* and *rpl-31* RNAi respectively at two hour intervals. Protein knockdown of reconstituted RPL split-GFP was then measured at the time of invasion. The optimal time for maximum knockdown for both *rpl-4* and *rpl-31* was 26.5 hours on RNAi (plating started at beginning of L2 larval stage), when RPL-31::GFP11 levels were reduced 20% compared to empty vector control (n ≥ 10 P6.p 2-cell ACs) and RPL-4::GFP11 levels were reduced 50% (n ≥ 10 P6.p 2-cell ACs).

### Image Quantification and Statistical Analysis

All fluorescence intensity measurements were performed in Fiji 1.53f (Schneider and Eliceiri, 2012). For comparison of mean fluorescence intensity or ER enrichment, either a pilot trial was conducted for each experiment, and a priori power analysis was performed to determine the appropriate sample size (α: 0.05, β: 0.80) or a *post-hoc* power analysis was conducted to determine the statistical power of the collected sample size (α: 0.05, β>0.80) using GPower 3.1. Statistical analyses were conducted and all graphs were generated in GraphPad Prism (Version 7). Statistical analyses comparing the percent defective invasion in treatment conditions to control were conducted using Fisher’s Exact 2x2 test. Comparisons of mean intensity or ER enrichment were done using either an unpaired two-tailed t-test or an unpaired two-tailed t-test with Welch’s correction when variances were different between the two groups based on an F-test. Comparisons of three or more conditions were made using a one-way ANOVA followed by a *post-hoc* Tukey’s test. Figure legends indicate sample sizes, statistical tests, and p-values. Experiments were not randomized, and investigators were not blinded during analysis. All figures were assembled in Adobe Illustrator Version 26.4.

