## Supplemental Figures & Tables for "The *C. elegans* Anchor Cell Transcriptome: Ribosome Biogenesis Drives Cell Invasion through Basement Membrane"

**SUPPLEMENTAL INFORMATION**

**Supplemental Tables**

**Table S1. Anchor cell (AC) transcriptome compared with mid L3 larval stage whole-body (WB) gene expression.**

Three replicates of AC and WB gene expression profiles are shown. Genes are ordered based upon log2 fold change values (the log2 transformed ratio of the average of the 3 AC replicates normalized reads over the average of the 3 WB replicates normalized reads). Genes listed were considered to be part of the AC transcriptome if they had at least 10 reads in one of the three AC replicates.

**Table S2. AC transcripts with significantly enriched expression versus mid L3 larval stage WB expression.**

Three replicates of AC and WB gene expression profiles are shown. Genes are ordered based upon log2 fold change values (the log2 transformed ratio of the average of the 3 AC replicates normalized reads over the average of the 3 WB replicates normalized reads). A list of 1,502 AC genes with log2 fold change ≥ 1 compared with WB gene expression and an adjusted p-value < 0.1.

**Table S3. Analysis of transgenic and genome edited reporters of genes present in the AC transcriptome.**

Complete list of genes previously annotated and newly identified as having AC expression based on visual analysis of transgenic or endogenously tagged strains. Log2 fold change is shown for each gene (the log2 transformed ratio of the average of the 3 AC replicates normalized reads over the average of the 3 WB replicates normalized reads). Sources are listed by PubMed reference number (PMID).

**Table S4. Previously tagged genes visually examined for AC expression.**

A list of genes that were present at various levels in the AC transcriptome and examined for expression in the AC at the P6.p 2-cell stage.  Expression was compared to other uterine and vulval cells using fluorescence microscopy, see Figures S2 and S3. Sources are listed by PubMed reference number (PMID).

**Table S5. A focused RNAi screen of AC enriched genes encoding secreted, transmembrane, cytoskeletal and transcription factor proteins.**

A focused RNAi screen of AC enriched genes (versus WB gene expression) encoding transmembrane or secreted proteins, cytoskeleton component and regulators, and transcription factors. AC invasion was scored by assessing the presence of a basement membrane (BM) breach using *lam-1p*::LAM-1::mCherry.

**Table S6. AC transcriptome enriched translational regulators.**

Genes encoding translational regulators enriched within the AC transcriptome versus WB gene expression. Genes were identified using Database for Annotation, Visualization and Integrated Discovery (DAVID) and WormBase ParaSite Biomart was used to identify human orthologs.

**Table S7. Genetic mutants examined for AC invasion defects.**

A list of genetic mutants examined for AC invasion defects. AC invasion was scored by DIC microscopy and assessing the presence or break in the phase-dense line under the AC, which corresponds to the BM.

**Table S8. Strains used in this study.**

**Table S9. Oligonucleotide sequences used in genome edited strains and RNAi constructs.**

**Table S3: Analysis of transgenic and genome edited reporters of genes in the AC transcriptome.**

| Genes previously identified as expressed in the AC | | | |
|---|---|---|---|
| **Gene** | **Sequence ID** | **Source (PMID)** | **log2 fold change** |
| *nhr-67* | C08F8.8 | 26506306 | 6.89 |
| *egl-43* | R53.3 | 20442418 | 5.15 |
| *hlh-2* | M05B5.5 | 21784067 | 4.93 |
| *lag-2* | Y73C8B.4 | 21784067 | 4.71 |
| *exc-6* | F58B6.2 | 26506306 | 4.26 |
| *zmp-6* | T21D11.1 | 26506306 | 3.9 |
| *lin-29* | W03C9.4 | 27661254 | 3.37 |
| *cdh-3* | ZK112.7 | 15960981 | 2.58 |
| *fos-1* | F29G9.4 | 15960981 | 2.39 |
| *unc-34* | Y50D4C.1 | 20442418 | 2.08 |
| *mig-2* | C35C5.4 | 19098902 | 2.08 |
| *lin-3* | F36H1.4 | 21784067 | 1.98 |
| *guk-1* | T03F1.8 | 20442418 | 1.94 |
| *tag-52* | C02F12.4 | 19540360 | 1.84 |
| *ephx-1* | K07D4.7 | 19540360 | 1.77 |
| *snap-29* | K02D10.5 | 29161591 | 1.72 |
| *gdi-1* | Y57G11C.10 | 26765257 | 1.6 |
| *egl-17* | F38G1.2 | 21784067 | 1.24 |
| *lit-1* | W06F12.1 | 20442418 | 1.23 |
| *cdc-37* | W08F4.8 | 20442418 | 1.22 |
| *mig-10* | F10E9.6 | 24553288 | 1.14 |
| *mig-6* | C37C3.6 | 21784067 | 1.1 |
| *hda-1* | C53A5.3 | 20442418 | 1.08 |
| *unc-115* | F09B9.2 | 24553288 | 1.04 |
| *cgef-1* | C14A11.3 | 19540360 | 1.03 |
| *unc-60* | C38C3.5 | 24662568 | 1.02 |
| *ant-1.1* | T27E9.1 | 30686527 | 0.94 |
| *unc-73* | F55C7.7 | 19540360 | 0.9 |
| *arx-2* | K07C5.1 | 30686527 | 0.88 |
| *cdc-42* | R07G3.1 | 26765257 | 0.73 |
| *mep-1* | M04B2.1 | 20442418 | 0.66 |
| *cct-7* | T10B5.5 | 20442418 | 0.58 |
| *ced-10* | C09G12.8 | 19098902 | 0.55 |
| *vab-10* | ZK1151.1 | 25443298 | 0.53 |
| *par-3* | F54E7.3 | 19098902 | 0.39 |
| *pix-1* | K11E4.4 | 19540360 | 0.34 |
| *F22G12.5* | F22G12.5 | 19540360 | 0.31 |
| *cacn-1* | W03H9.4 | 20442418 | 0.19 |
| *tag-77* | C28C12.10 | 19540360 | 0.13 |
| *ina-1* | F54G8.3 | 19686680 | -0.03 |
| *ced-5* | C02F4.1 | 19540360 | -0.04 |

| | | | |
|---|---|---|---|
| *ect-2* | T19E10.1 | 19540360 | -0.12 |
| *unc-40* | T19B4.7 | 19686680 | -0.13 |
| *zmp-1* | EGAP1.3 | 15960981 | -0.27 |
| *wsp-1* | C07G1.4 | 26765257 | -0.56 |
| *mdt-28* | F28F8.5 | 20442418 | -0.58 |
| *exc-5* | C33D9.1 | 19540360 | -1.03 |
| *him-4* | F15G9.4 | 15960981 | -1.06 |
| *pitr-1* | C48A7.2 | 20442418 | -1.15 |
| *pat-3* | ZK1058.2 | 19686680 | -1.17 |
| *zmp-3* * | C31B8.8 | 26506306 | -1.53 |
| *trd-1* | T20B12.1 | 20442418 | -2.23 |
| **Genes identified as expressed in the AC in this study through endogenous tagging** | | | |
| **Gene** | **Sequence ID** | **Source (PMID)** | **log2 fold change** |
| *dmd-3* | Y43F8C.10 | Current Study | 5.52 |
| *rap-1* | C27B7.8 | Current Study | 2.9 |
| *tct-1* | F25H2.11 | Current Study | 1.94 |
| *snb-1* | T10H9.4 | Current Study | 1.79 |
| *ebp-2* | VW02B12L.3 | Current Study | 1.69 |
| *rab-11.1* | F53G12.1 | Current Study | 1.51 |
| *eif-1.A* | H06H21.3 | Current Study | 0.9 |
| *sdn-1* | F57C7.3 | Current Study | 0.4 |
| *ten-1* | R13F6.4 | Current Study | 0.11 |
| *ptp-3* | C09D8.1 | Current Study | -0.16 |
| **Gene whose expression was not detected in the AC in this study through endogenous tagging** | | | |
| **Gene** | **Sequence ID** | **Source (PMID)** | **log2 fold change** |
| *air-2* | B0207.4 | Current Study | -1.33 |

* denotes having less than 10 reads in AC libraries.

**Table S4: Previously tagged genes visually examined for AC expression.**

| Gene | Sequence ID | Source (PMID) | Notes | Fluorophore tagging method |
|---|---|---|---|---|
| *nhr-67* | C08F8.8 | 31806663 | nuclear hormone receptor Tailless/TLX | endogenous tag |
| *fos-1* | F29G9.4 | 31806663 | basic leucine zipper transcription factor Fos | endogenous tag |
| *hlh-2* | M05B5.5 | 31806663 | basic helix-loop-helix transcription facotr E/Daughterless | endogenous tag |
| *egl-43* | R53.3 | 31806663 | zinc-finger transcription factor EVI1/MEL | endogenous tag |
| *dmd-3* | Y43F8C.10 | 30599092 | doublesex | endogenous tag |
| *ina-1* | F54G8.3 | 32585132 | alpha integrin | endogenous tag |
| *pat-3* | ZK1058.2 | 32585132 | beta integrin | endogenous tag |
| *ptp-3* | C09D8.1 | 32585132 | LAR-type receptor phosphotyrosine-phosphatases | endogenous tag |
| *sdn-1* | F57C7.3 | 32585132 | syndecan | endogenous tag |
| *lag-2* | Y73C8B.4 | 30799241 | Notch receptor ligand | endogenous tag |
| *ebp-2* | VW02B12L.3 | 30080857 | homolog of microtubule tip tracking protein EBP-1 | endogenous tag |
| *arx-2* | K07C5.1 | 27780040 | Arp2/3 subunit | endogenous tag |
| *rap-1* | C27B7.8 | 28829947 | small GTPase | endogenous tag |
| *guk-1* | T03F1.8 | 20442418 | guanylate kinase | translational reporter |
| *air-2* | B0207.4 | 34014923 | Aurora B | endogenous tag |
| *zmp-1* | EGAP1.3 | 30686527 | GPI anchored MMP | endogenous tag |

**Table S5: A focused RNAi screen of AC enriched genes encoding secreted, transmembrane, cytoskeletal and transcription factor proteins.**

| | RNAi treatment | Sequence ID | % ACs with incomplete invasion[a] | p value[b] | *n*[c] |
|---|---|---|---|---|---|
| Negative Control | | | | | |
| 1 | L4440 (Empty Vector) | N/A | 0% | N/A | 60 |
| Positive Control | | | | | |
| 2 | *fos-1* | F29G9.4 | 78% | <0.0001 | 60 |
| Enriched AC transcriptome genes | | | | | |
| 3 | *tct-1* | F25H2.11 | 34% | 0.0005 | 12 |
| 4 | *fmi-1* | F15B9.7 | 19% | 0.004 | 21 |
| 5 | *mom-1* | T07H6.2 | 20% | 0.007 | 15 |
| 6 | *hhat-2* | Y57G11C.17 | 16% | 0.01 | 26 |
| 7 | *K07C11.7* | K07C11.7 | 17% | 0.03 | 12 |
| 8 | *dab-1* | M110.5 | 13% | 0.04 | 15 |
| 9 | *col-155* | F55C10.3 | 13% | 0.04 | 15 |
| 10 | *cyn-5* | F31C3.1 | 13% | 0.04 | 15 |
| 11 | *snb-1* | T10H9.4 | 14% | 0.04 | 15 |
| 12 | *wrt-6* | ZK377.1 | 14% | 0.04 | 15 |
| 13 | *tmed-3* | F57B10.5 | 14% | 0.04 | 15 |
| 14 | *paqr-3* | Y67A10A.8 | 12% | 0.04 | 16 |
| 15 | *mlc-4* | C56G7.1 | 12% | 0.05 | 18 |
| 16 | *ccr-4* | ZC518.3 | 10% | ns | 19 |
| 17 | *unc-116* | R05D3.7 | 9% | ns | 22 |
| 18 | *chc-1* | T20G5.1 | 8% | ns | 23 |
| 19 | *tbb-2* | C36E8.5 | 17% | ns | 6 |
| 20 | *sec-61.G* | F32D8.6 | 10% | ns | 10 |
| 21 | *pdcd-2* | R07E5.10 | 8% | ns | 12 |
| 22 | *hint-1* | F21C3.3 | 8% | ns | 13 |
| 23 | *pqn-25* | D1044.3 | 8% | ns | 13 |
| 24 | *sbsp-1* | R17.3 | 7% | ns | 14 |
| 25 | *syd-2* | F59F5.6 | 7% | ns | 15 |
| 26 | *dmd-3* | Y43F8C.10 | 7% | ns | 15 |
| 27 | *smo-1* | K12C11.2 | 7% | ns | 15 |
| 28 | *ben-1* | C54C6.2 | 7% | ns | 15 |
| 29 | *nphp-4* | R13H4.1 | 7% | ns | 15 |
| 30 | *lst-2* | R160.7 | 7% | ns | 15 |
| 31 | *szy-2* | Y32H12A.4 | 7% | ns | 15 |
| 32 | *dpy-5* | F27C1.8 | 7% | ns | 15 |
| 33 | *M03A1.3* | M03A1.3 | 7% | ns | 15 |
| 34 | *grk-2* | W02B3.2 | 6% | ns | 16 |
| 35 | *ptc-3* | Y110A2AL.8 | 6% | ns | 17 |

| | | | | | |
|---|---|---|---|---|---|
| 36 | *F55A11.1* | F55A11.1 | 6% | ns | 18 |
| 37 | *par-5* | M117.2 | 5% | ns | 19 |
| 38 | *lgg-3* | B0336.8 | 4% | ns | 25 |
| 39 | *afd-1* | W03F11.6 | 0% | ns | 15 |
| 40 | *stam-1* | C34G6.7 | 0% | ns | 15 |
| 41 | *pept-3* | F56F4.5 | 0% | ns | 15 |
| 42 | *suro-1* | R11A5.7 | 0% | ns | 15 |
| 43 | *snx-3* | W06D4.5 | 0% | ns | 12 |
| 44 | *ztf-11* | F52F12.6 | 0% | ns | 15 |
| 45 | *rab-3* | C18A3.6 | 0% | ns | 15 |
| 46 | *sqv-8* | ZK1307.5 | 0% | ns | 15 |
| 47 | *unc-53* | F45E10.1 | 0% | ns | 11 |
| 48 | *exc-6* | F58B6.2 | 0% | ns | 15 |
| 49 | *rnf-5* | C16C10.7 | 0% | ns | 15 |
| 50 | *frm-2* | T04C9.6 | 0% | ns | 15 |
| 51 | *ykt-6* | B0361.10 | 0% | ns | 17 |
| 52 | *tbb-1* | K01G5.7 | 0% | ns | 15 |
| 53 | *cccp-1* | Y49E10.23 | 0% | ns | 19 |
| 54 | *exc-9* | F20D12.5 | 0% | ns | 15 |
| 55 | *rap-1* | C27B7.8 | 0% | ns | 15 |
| 56 | *nas-7* | C07D10.4 | 0% | ns | 30 |
| 57 | *pcm-1* | C10F3.5 | 0% | ns | 15 |
| 58 | *ptr-1* | C24B5.3 | 0% | ns | 15 |
| 59 | *mics-1* | T21C9.1 | 0% | ns | 15 |
| 60 | *max-1* | C34B4.1 | 0% | ns | 15 |
| 61 | *fmo-4* | F53F4.5 | 0% | ns | 15 |
| 62 | *hif-1* | F38A6.3 | 0% | ns | 15 |
| 63 | *sec-22* | F55A4.1 | 0% | ns | 11 |
| 64 | *snx-1* | C05D9.1 | 0% | ns | 23 |
| 65 | *sax-1* | R11G1.4 | 0% | ns | 15 |
| 66 | *haly-1* | F47B10.2 | 0% | ns | 15 |
| 67 | *ptrn-1* | F35B3.5 | 0% | ns | 15 |
| 68 | *dkf-1* | W09C5.5 | 0% | ns | 15 |
| 69 | *dgk-1* | C09E10.2 | 0% | ns | 15 |
| 70 | *dnc-2* | C28H8.12 | 0% | ns | 15 |
| 71 | *bar-1* | C54D1.6 | 0% | ns | 12 |
| 72 | *ift-81* | F32A6.2 | 0% | ns | 15 |
| 73 | *mbl-1* | K02H8.1 | 0% | ns | 15 |
| 74 | *hst-3.2* | F52B10.2 | 0% | ns | 15 |
| 75 | *ric-8* | Y69A2AR.2 | 0% | ns | 14 |
| 76 | *fkb-3* | C05C8.3 | 0% | ns | 15 |
| 77 | *sbt-1* | T03D8.3 | 0% | ns | 24 |
| 78 | *nhx-5* | F57C7.2 | 0% | ns | 14 |
| 79 | *efn-2* | C43F9.8 | 0% | ns | 23 |
| 80 | *txdc-12.1* | Y57A10A.23 | 0% | ns | 12 |
| 81 | *clk-1* | ZC395.2 | 0% | ns | 15 |
| 82 | *hlh-12* | C28C12.8 | 0% | ns | 15 |

| | | | | | |
|---|---|---|---|---|---|
| 83 | *tbcb-1* | F53F4.3 | 0% | ns | 12 |
| 84 | *dnc-2* | C28H8.12 | 0% | ns | 15 |

[a] A *rrf-3* RNAi sensitized strain (see Methods) was used for scoring invasion with an AC plasma membrane marker (*cdh-3p*::GFP::CAAX) and BM marker (*lam-1p*::LAM-1::mCherry). All animals were scored for invasion at the VPC P6.p 4-cell stage (see Figure 1A).

[b] Fischer Exact 2x2 Tests were used. RNAi knockdown effects on AC invasion were compared to L4440 (empty vector) control for statistical analysis.

[c] Number of animals observed for each condition.

**Table S6: AC transcriptome enriched translational regulators**

| Gene | Sequence ID | Human Orthologs | Ensembl Stable ID |
|---|---|---|---|
| *asd-2* | T21G5.5 | QKI | ENSG00000112531 |
| *eif-1* | T27F7.3 | EIF1B | ENSG00000114784 |
| *ife-2* | R04A9.4 | EIF4E2 | ENSG00000135930 |
| *ife-2* | R04A9.4 | EIF4E3 | ENSG00000163412 |
| *ife-4* | C05D9.5 | ENSG00000386996 | ENSG00000386996 |
| *mrps-11* | W04D2.5 | MRPS11 | ENSG00000181991 |
| *rpl-5* | F54C9.5 | RPL5 | ENSG00000122406 |
| *rpl-19* | C09D4.5 | RPL19 | ENSG00000108298 |
| *rpl-21* | C14B9.7 | RPL21 | ENSG00000122026 |
| *rpl-27* | C53H9.1 | RPL27 | ENSG00000131469 |
| *rpl-29* | B0513.3 | RPL29 | ENSG00000162244 |
| *rpl-30* | Y106G6H.3 | RPL30 | ENSG00000156482 |
| *rpl-31* | W09C5.6 | RPL31 | ENSG00000071082 |
| *rpl-33* | F10E7.7 | RPL35A | ENSG00000182899 |
| *rpl-34* | C42C1.14 | RPL34 | ENSG00000109475 |
| *rpl-37* | C54C6.1 | RPL37 | ENSG00000274242 |
| *rpl-38* | C06B8.8 | RPL38 | ENSG00000172809 |
| *rpl-39* | C26F1.9 | RPL39 | ENSG00000355315 |
| *rps-10* | D1007.6 | RPS10 | ENSG00000124614 |
| *rps-22* | F53A3.3 | RPS15A | ENSG00000134419 |
| *rps-27* | F56E10.4 | RPS27L | ENSG00000331019 |
| *snr-1* | Y116A8C.42 | SNRPD3 | ENSG00000100028 |
| *snr-4* | C52E4.3 | SNRPD2 | ENSG00000125743 |
| *snr-5* | ZK652.1 | SNRPF | ENSG00000139343 |
| *snr-6* | Y49E10.15 | SNRPE | ENSG00000182004 |
| *snrp-27* | R05D11.7 | SNRNP27 | ENSG00000124380 |

**Table S7: Genetic mutants examined for AC invasion defects.**

| | Identifier | Genotype | % ACs with incomplete invasion[a] | p value[b] | *n*[c] |
|---|---|---|---|---|---|
| 1 | N2 | Wild-type | 0% | n/a | 50 |
| 2 | RB925 | *ire-1(ok799)* II | 25% | 0.001 | 20 |
| 3 | MC817 | *ddx-52(gc51)* I | 16% | 0.01 | 25 |
| 4 | FK312 | *sma-5(n678)* X | 0% | ns | 25 |
| 5 | NK2694 | *qy110(rpl-31-GFP11)* I, *bmd15(eft-3p::GFP1-10::unc54 3'UTR-[let858 terminator]-myo-2p::mCherry-3xHA tbb-2 3'UTR')* I | 0% | n/a | 41 |
| 6 | NK2902 | *qyIs463(lin-29p::ZIF-1::SL2::mCh)* , *rpl-31(qy189[rpl-31::ZF1::GFP11])* I, *bmd15(eef-1A.1p::GFP1-10::unc-54 3'UTR; myo-2p::mCherry tbb-2 3'UTR)* I; *zif-1(gk117)* III | 45% | <0.0001 | 40 |

[a] All animals were scored for invasion at the VPC P6.p 4-cell stage (see Figure 1A).

[b] Fischer Exact 2x2 Tests were used. Genetic mutants compared to N2 for statistical analysis. NK2902 was compared to NK2694 for statistical analysis.

[c] Number of animals observed for each condition.

**Table S8: Strains used in study.**

| Genotype | Description | Source (PMID) | Identifier |
|---|---|---|---|
| *unc-119(ed3)* III | *C. elegans* unc-119 integrated and extrachromosomal array injection strain | Caenorhabditis Genetics Center | WB Strain |
| *qyIs362(lin-29p::GFP); qyIs17(zmp-1p::mCherry)* II | AC specific cytoplasmic GFP and mCherry marked AC strain | 20442418 | NK1382 |
| ot932(dmd-3::GFP::3xFLAG) V; him-8 (e1489) IV | DMD-3::GFP endogenous tag | 30599092 | OH15733 |
| *qy161(tct-1::mNG)* I | TCT-1::mNG endogenous tag | This study | NK2800 |
| *cp151(mNG-C1::3xFLAG::rap-1)* IV | mNG::RAP-1 endogenous tag | 28829947 | LP399 |
| *qy56(ten-1::mNG)* III | TEN-1::mNG endogenous tag | 32585132 | NK2502 |
| *ie31(degron::GFP::air-2)* I | GFP::AIR-2 endogenous tag | 34014923 | CA1216 |
| *syb509(nhr-67::GFP)* IV | NHR-67::GFP endogenous tag | 31806663 | PHX509 |
| *bmd88(LoxP::GFP::egl-43)* II | GFP::EGL-43 endogenous tag | 31806663 | DQM300 |
| *bmd90(LoxP::GFP::hlh-2)* I | GFP::HLH-2 endogenous tag | 31806663 | DQM311 |
| *bmd138(LoxP::GFP::FOS-1)* V | GFP::FOS-1 endogenous tag | 31806663 | DQM497 |
| *qyIs49(guk-1p::GUK-1::YFP)* V; *unc-119(+)* | *guk-1* translational marker | 20442418 | NK370 |
| *qy164(snb-1::mNG)* V | SNB-1::mNG endogenous tag | This study | NK2827 |
| *qy190(mNG::rab-11.1)* I | mNG::RAB-11.1 endogenous tag | This study | NK2904 |
| *cp226(lin-3::mNG-C1::3xFLAG)* IV | LIN-3::mNG endogenous tag | This study | LP511 |
| *cas607(arx-2::GFP)* V | ARX-2::GFP endogenous tag | 27780040 | NK2144 |
| *qy90(eif-1.A::mNG)* IV | EIF-1.A::mNG endogenous tag | This study | NK2621 |
| *qy17(zmp-1::mNG)* III | ZMP-1::mNG endogenous tag | 30686527 | NK2144 |
| *qy29(sdn-1::mNG)* X | SDN-1::mNG endogenous tag | 32585132 | NK2413 |
| *qy23(ina-1::mNG)* III | INA-1::mNG endogenous tag | 31387941 | NK2324 |
| *qy47(ptp-3::mNG)* II | PTP-3::mNG endogenous tag | 32585132 | NK2477 |
| *qy36(pat-3::mNG)* III | PAT-3::mNG endogenous tag | 31387941 | NK2436 |
| *qyIs287(myo-3p::mCherry); cp193(lag-2::mNG::3xFLAG)* V | LAG-2::mNG endogenous tag | 30799241 | NK2376 |
| *qy110(rpl-31::gfp11)* I, *bmd15(eef-1A.1p::GFP1-10::unc-54 3'UTR; myo-2p::mCherry tbb-2 3'UTR)* I | RPL-31::GFP11 endogenous tag with single copy insertion of GFP1-10 driven by ubiquitous promoter | This study | NK2694 |
| *qy128(rpl-4::gfp11)* I, *bmd15(eef-1A.1p::GFP1-10::unc-54 3'UTR; myo-2p::mCherry tbb-2 3'UTR)* I | RPL-4::GFP11 endogenous tag with single copy insertion of GFP1-10 driven by ubiquitous promoter | This study | NK2730 |

| | | | |
|---|---|---|---|
| *qyIs463(lin-29p::ZIF-1::SL2::mCherry); qy189(rpl-31::ZF1::GFP11)* I; *bmd15(eef-1A.1p::GFP1-10::unc-54 3'UTR; myo-2p::mCherry tbb-2 3'UTR); zif-1(gk117)* III | RPL-31::ZF1::GFP11 endogenous tag with single copy insertion of GFP1-10 driven by ubiquitous promoter, with AC specific rescue of *zif-1*, in *zif-1* mutant background | This study | NK2902 |
| *wow47(ebp-2::GFP::3xFLAG)* II | EBP-2::GFP endogenous tag | This study | SBW292 |
| *shy61(sec-61.B::GFP11x2)* IV; *bmd15(eef-1A.1p::GFP1-10::unc-54 3'UTR; myo-2p::mCherry tbb-2 3'UTR)* I | SEC-61.B::GFP11 endogenous tag. | This study | NK2789 |
| *wgIs506(xbp-1::TY1::eGFP::3xFLAG)* | *xbp-1* translational marker. | 16990816 | OP506 |
| *qy97(elo-1::mNG)* IV | ELO-1::mNG endogenous tag. | This study | NK2633 |
| *rrf-3 (pk1426)* II; *qyIs127(lam-1p::lam-1::mCherry)* V; *qyIs166 (cdh-3p::GFP::CAAX)* X; *unc-119(+)* | Sensitized RNAi strain containing AC and BM markers for scoring invasion | 29161591 | NK1339 |
| *rrf-3 (pk1426)* II; *qyIs10(lam-1p::lam-1::GFP)* IV; *rde-1(ne219)* V; *qyIs24(cdh-3p::mCherry::PLCδPH); qyIs102(fos-1p::rde-1, myo-2p::GFP); unc-119(+)* | AC specific RNAi strain containing AC and BM markers for scoring invasion | 25443298 | NK1316 |
| *knuSi221(fib-1p::FIB-1::eGFP::fib-1 3'UTR)* | *fib-1* translational marker | 25298536 | COP262 |
| *qy206(dgn-1::GFP)* X | DGN-1::GFP endogenous tag | This study | NK2009 |
| *sma-5(n678)* X; *qyIs23(cdh-3p::mCherry::PLCδPH)* II | *sma-5* mutant with AC marker | This study | NK2963 |
| *qyEx603(lin-29p::AMAN-2(aa1-84)::mScarlet); unc-119(+)* | Strain containing golgi marker | This study | NK2880 |
| *qyIs66(cdh-3p::unc-40::GFP); rrf-3(pk1426)* II; *unc-119(+)* | AC specific UNC-40::GFP translational reporter in RNAi sensitized background | 23751497 | NK455 |
| *qyIs57(cdh-3p::mCherry::moesinABD); rrf-3(pk1426)* II; *unc-119(+)* | AC specific F-actin reporter in RNAi sensitized background | This study | NK2960 |
| *qyIs550(zmp-1p::MLS::GFP); rrf-3(pk1426)* II | AC specific mitochondria reporter in RNAi sensitized background | This study | NK2961 |
| *qy17(zmp-1::mNG::GPI)* III; *rrf-3(pk1426)* II | ZMP-1::mNG endogenous tag in RNAi sensitized background | This study | NK2962 |
| *qy65(fgt-1::mNG)* II | FGT-1::mNG endogenous tag | 35316617 | NK2540 |
| *qy126(nifk-1::mNG)* III; *zmp-1(cg115)* III | NIFK-1::mNG endogenous tag in zmp-1 mutant strain | This study | NK2964 |
| *qy121(eef-1A.1p::GFP)* I | Single copy insertion of *eef-1A.1p*::GFP | This study | NK2790 |

| | | | |
|---|---|---|---|
| *bmd243(LoxN,cdh-3p::TIR1::F2A::DHB::2xmTurquoise2)* I; *ljf3(unc-34::mNG[C1]^3xFlag::AID)* V; *qy41(lam-2::mKate2)* X | UNC-34::mNG endogenous tag | This study | DQM1152 |
| *ddx-52 (gc51)* I | *ddx-52* mutant | 33157031 | MC817 |
| *ire-1(ok799)* II | *ire-1* mutant | 23173093 | RB925 |
| *qyIs10(lam-1p::LAM-1::GFP)* IV; *qyIs24(cdh-3p::mCherry::PLCδPH); unc-119(+)* | Strain containing AC and BM markers | 19540360 | NK361 |
| *unc-119(tm4063)* III; *wgIs793(cep-1::TY1::eGFP::3xFLAG); unc-119(+)* | Strain containing p53 marker | 20174564 | OP793 |

**Table S9: Oligonucleotide sequences used in genome edited strains and RNAi constructs.**

| Oligonucleotide Sequence (5'->3') | Primer type | Amplicon | Template |
|---|---|---|---|
| Cas9 Short Guide Primers | | | |
| tcctattgcgagatgtctttacagctcagcagtaaagatgttttagagctagaaatagc | Forward | *tct-1* sgRNA | pDD122 |
| tcctattgcgagatgtctttacagctcagcagtaaagatgttttagagctagaaatagc | Forward | *rab-11.1* sgRNA | pDD122 |
| tcctattgcgagatgtctttaagtacacgacctttgtccgttttagagctagaaatagc | Forward | *snb-1* sgRNA | pDD122 |
| tcctattgcgagatgtcttcgtggtggcaagaacaagtagttttagagctagaaatagc | Forward | *eif-1.A* sgRNA | pDD122 |
| tcctattgcgagatgtcttcttttcgacagcagctggacgttttagagctagaaatagc | Forward | *nifk-1* sgRNA | pDD122 |
| tcctattgcgagatgtcttcgcggagaacatatgattgggttttagagctagaaatagc | Forward | *elo-1* sgRNA 1 | pDD122 |
| tcctattgcgagatgtcttctccgcggagaacatatgatgttttagagctagaaatagc | Forward | *elo-1* sgRNA 2 | pDD122 |
| tcctattgcgagatgtctttttcatgtgtgtcgaatcatgttttagagctagaaatagc | Forward | *lin-3* sgRNA | pDD122 |
| tcctattgcgagatgtcttgaatacttatttcttcacgtgttttagagctagaaatagc | Forward | *rpl-4* sgRNA 1 | pDD122 |
| tcctattgcgagatgtcttcgttggcctttgaagccttggttttagagctagaaatagc | Forward | *rpl-4* sgRNA 2 | pDD122 |
| tcctattgcgagatgtcttcaacgtcaacgttgacagcggttttagagctagaaatagc | Forward | *rpl-31* sgRNA | pDD122 |
| tcctattgcgagatgtcttgagctatataggctctatcgagttttagagctagaaatagc | Forward | *unc-34* sgRNA | pDD122 |
| tcctattgcgagatgtcttccacgtctcggacatgcttcatcgttttagagctagaaatagc | Forward | *dgn-1* sgRNA | pDD122 |
| Genotyping Primers | | | |
| gaatacaccattcacatccacg | Forward | *rpl-31* genotyping external | *rpl-31::GFP11* gDNA or *rpl-31::ZF1::GFP11* gDNA |
| cacgtggctgcaaatattcga | Reverse | *rpl-31* genotyping external | *rpl-31::GFP11* gDNA or *rpl-31::ZF1::GFP11* gDNA |

| | | | |
|---|---|---|---|
| caaccgtcaacacaagcagaag | Forward | *rpl-4* genotyping external | *rpl-4::GFP11* gDNA |
| gtttcatctctcgttcgtgc | Reverse | *rpl-4* genotyping external | *rpl-4::GFP11* gDNA |
| ccttgtccgaccatgtcgaa | Reverse | mNG genotyping | gDNA |
| tttgtatagttcatccatgccatgtg | Reverse | GFP genotyping | gDNA |
| ctccaaaagcagctactccaa | Forward | *nifk-1* genotyping external | *nifk-1::mNG* gDNA |
| aaggctgatgtcgatgcctt | Forward | *tct-1* genotyping external | *tct-1::mNG* gDNA |
| ataggcgattgattacctgga | Forward | *rab-11.1* genotyping external | *mNG::rab-11.1* gDNA |
| tagagatgcgcgtaggtcat | Forward | *snb-1* genotyping external | *snb-1::mNG* gDNA |
| ccaagtccgcagttacgtaacc | Forward | *eif-1a* genotyping external | *eif-1.A::mNG* gDNA |
| aaatggttcatgcgtaatcgtc | Forward | *elo-1* genotyping external | *elo-1::mNG* gDNA |
| ggattccggcccacttccttc | Forward | *dgn-1* genotyping external | *dgn-1::GFP* gDNA |
| ttcagaatcgtgtctcccttcgtggtttcg | Forward | *lin-3* genotyping external | *lin-3::mNG* gDNA |
| tggtttagaagtagccacgagttg | Forward | *sma-5 (n678)* genotyping external | *sma-5 (n678)* ; qyIs23 gDNA |
| gaagcaagatgtccgatcagcg | Reverse | *sma-5 (n678)* genotyping external | *sma-5 (n678)* ; qyIs23 gDNA |
| cacgcatccatcctttcaccag | Reverse | *sma-5 (n678)* genotyping nested | *sma-5 (n678)* ; qyIs23 gDNA |

| | | | |
|---|---|---|---|
| gatgctaagctcattggcag | Forward | *rrf-3 (pk1426)* genotyping, external | *rrf-3 (pk1426)* crossed strain DNA |
| ccaatgcatcttccagcctt | Reverse | *rrf-3 (pk1426)* genotyping, external | *rrf-3 (pk1426)* crossed strain DNA |
| tgaacgcggtaactacatcg | Reverse | *rrf-3 (pk1426)* genotyping, nested | *rrf-3 (pk1426)* crossed strain DNA |
| cttatgggcgtctcgtgt | Forward | *zif-1 (gk117)* genotyping | *zif-1 (gk117)* crossed stain DNA |
| tcatcatccggtcctgc | Reverse | *zif-1 (gk117)* genotyping | *zif-1 (gk117)* crossed strain DNA |
| Transgenic Strain Primers | | | |
| aagggcccatgggaaaacgcaatttcta | Forward | aman-2(aa1-84) with APAI overhang | pHD93 |
| aagtcgacttctttttcttcatcaaaat | Reverse | aman-2(aa1-84) with SALI overhang | pHD93 |
| atcggtagattttgatgaagaaaaagaagtctcc aagggagaggccgtcatcaagg | Forward | aman-2(aa1-84) with mScarlet overhang | *pAP088-lin-29p::mScarlet* |
| gaaattgcgttttcccatgcatgctgcgttgaaga agttggcttga | Reverse | aman-2(aa1-84) with lin-29 promoter overhang | *pAP088-lin-29p::mScarlet* |
| cgacggccagtaaagctagctttattgtcaacttc cattg | Forward | eef-1A.1 promoter with pCFJ352 overhang | pDD162 |
| cgacggccagtaaagctagctttattgtcaacttc cattg | Reverse | eef-1A.1 promoter with GFP overhang | pDD162 |
| tagcgaccggcgctcagttggcggccgcctattt gtatagttcatccatgcc | Reverse | GFP with pCFJ352 overhang | *pBS-cdh-3:: GFP* |
| RNAi Primers | | | |
| agatctgatatcatcgatgaattcgagcatggctt atctgggtgtcgata | Forward | pro-1 RNAi with L4440 backbone overhang | N2 gDNA |

| | | | |
|---|---|---|---|
| actcactatagggcgaattgggtaccttcgctatcaacaatctccttgg | Reverse | pro-1 RNAi with L4440 backbone overhang | N2 gDNA |
| atgaattcgagctccaccgcggtatgctgatctacaaggatat | Forward | tct-1 RNAi wtih T444T vector backbone overhang | N2 gDNA |
| ggtcgacggtatcgataagcttgagcacttctcctcgatgatgg | Reverse | tct-1 RNAi wtih T444T vector backbone overhang | N2 gDNA |

### Supplemental Figures

#### FIGURE S1

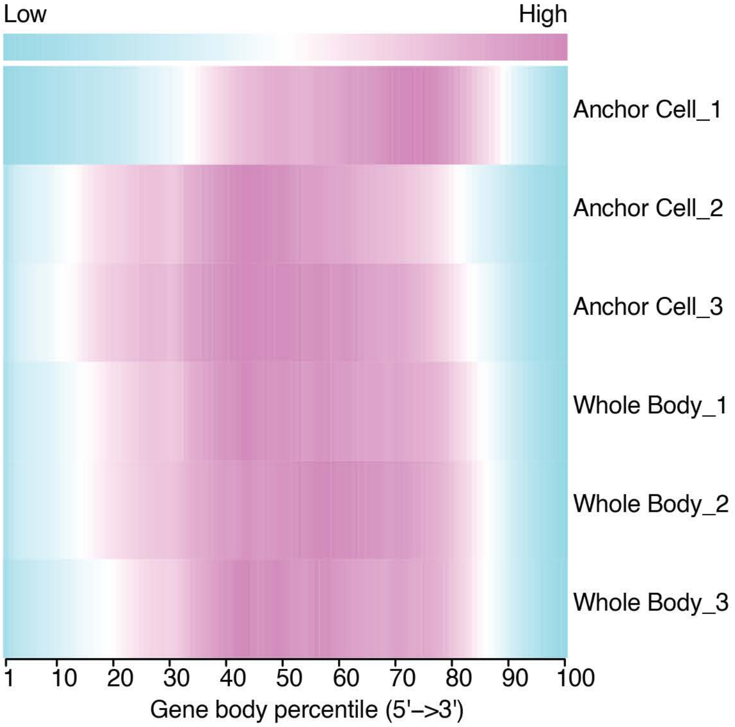

**Figure S1. Gene body coverage analysis of single-cell RNA sequencing.**

Heatmap of read coverage profiles over gene body to evaluate uniformity of 5’ to 3’ coverage (pink = high coverage, blue = low coverage).

FIGURE S2

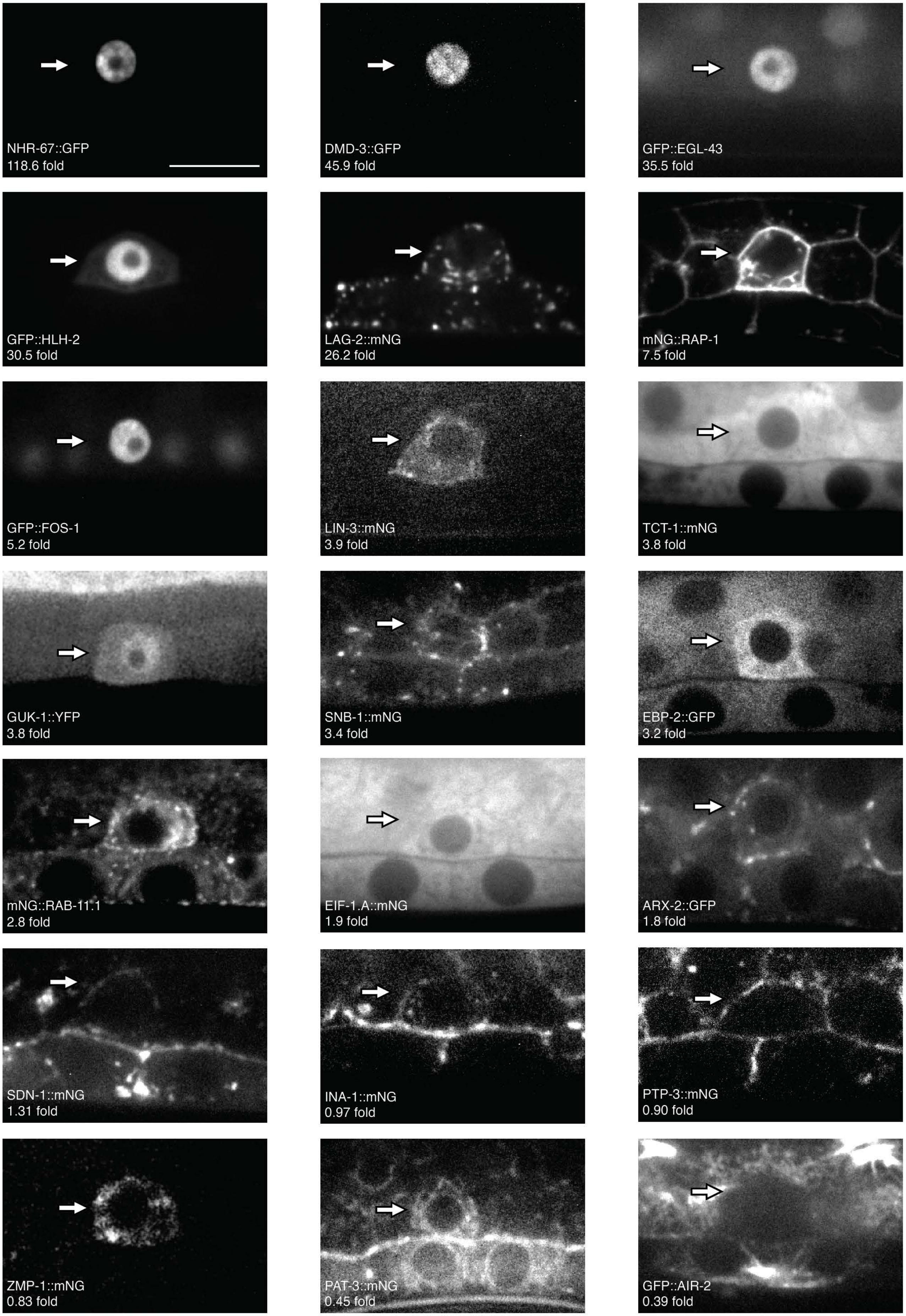
NHR-67::GFP
118.6 fold
DMD-3::GFP
45.9 fold
GFP::EGL-43
35.5 fold
GFP::HLH-2
30.5 fold
LAG-2::mNG
26.2 fold
mNG::RAP-1
7.5 fold
GFP::FOS-1
5.2 fold
LIN-3::mNG
3.9 fold
TCT-1::mNG
3.8 fold
GUK-1::YFP
3.8 fold
SNB-1::mNG
3.4 fold
EBP-2::GFP
3.2 fold
mNG::RAB-11.1
2.8 fold
EIF-1.A::mNG
1.9 fold
ARX-2::GFP
1.8 fold
SDN-1::mNG
1.31 fold
INA-1::mNG
0.97 fold
PTP-3::mNG
0.90 fold
ZMP-1::mNG
0.83 fold
PAT-3::mNG
0.45 fold
GFP::AIR-2
0.39 fold

**Figure S2. Levels of fluorescently tagged proteins of select AC transcriptome genes at the P6.p 2-cell AC.**

AC (arrows) protein expression of endogenous reporters generated through genome editing, except GUK-1::YFP which was generated as a translational reporter. Fold change of the AC compared to whole-body transcriptomes are shown (bottom left of each image). DMD-3, RAP-1, TCT-1, SNB-1, EBP-2, RAB-11.1, and AIR-2 are also shown in Figure 1, but included here for completeness. Scale bar: 5 µm.

### FIGURE S3

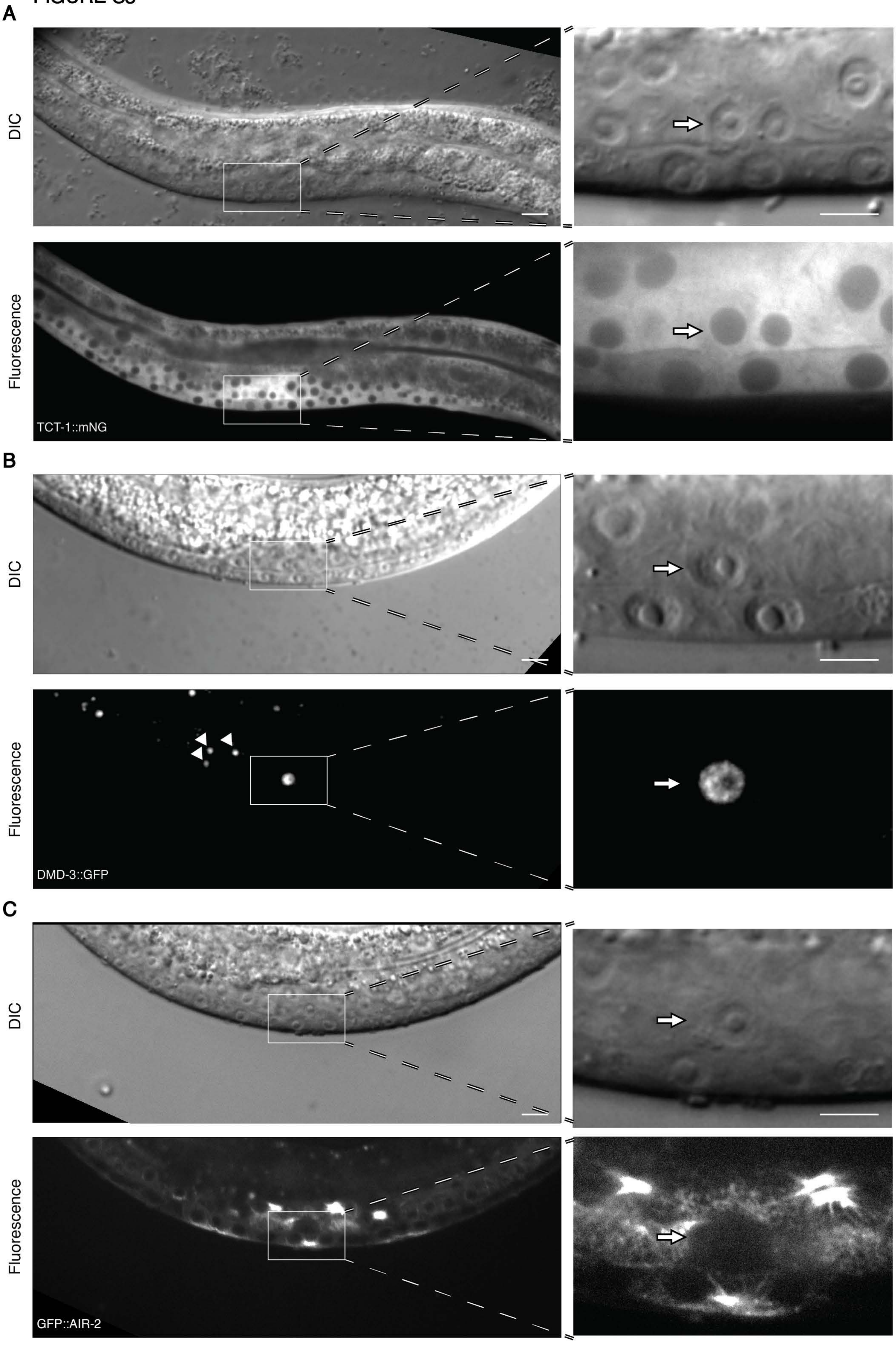
A
DIC
Fluorescence
TCT-1::mNG
B
DIC
Fluorescence
DMD-3::GFP
C
DIC
Fluorescence
GFP::AIR-2

**Figure S3. Levels and cellular distribution of endogenously tagged proteins of select AC enriched and underenriched genes.**

(A-C) Endogenous proteins in ACs (arrows) at the P6.p 2-cell stage when the AC transcriptome was generated. AC enrichment was based on comparison to WB gene expression at the same developmental stage. A central body view (left, 400x magnification) showing where the AC is located (box) is compared to a magnified view of the AC region (1000X magnification, right). Endogenous tagging of translational proteins generated from the highly enriched *tct-1* and *dmd-3* genes (A,B) and the underenriched *air-2* (C) gene. Scale bars: Left, 10 μm. Right, 5 μm.

FIGURE S4

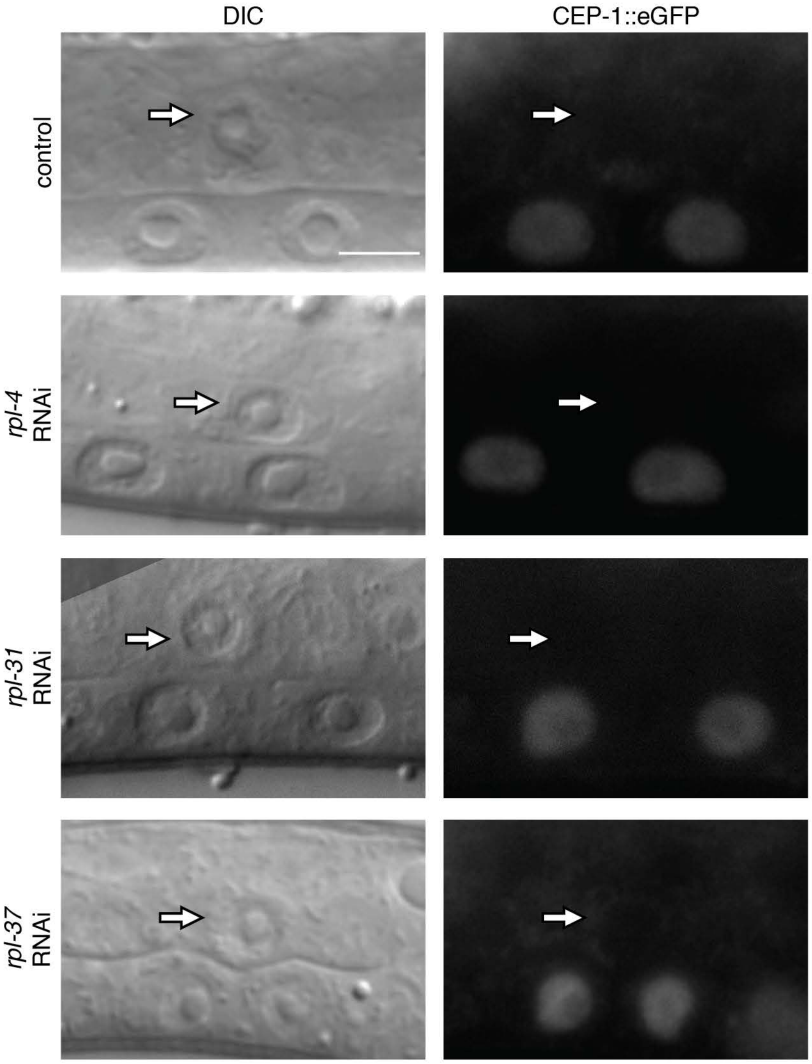

**Figure S4. CEP-1 (p53) is not stabilized/activated in the AC after RNAi mediated reduction of large ribosomal subunit proteins.**

Examples of ACs (left, DIC; right, CEP-1::eGFP (p53); arrow) at the P6.p 2-cell stage after treatment with control RNAi and after RNAi mediated knockdown of *rpl-4, rpl-31,* and *rpl-37*. CEP-1::eGFP was not detected in the AC after any treatment (n = 10 for each). Scale bars: 5 μm.

FIGURE S5

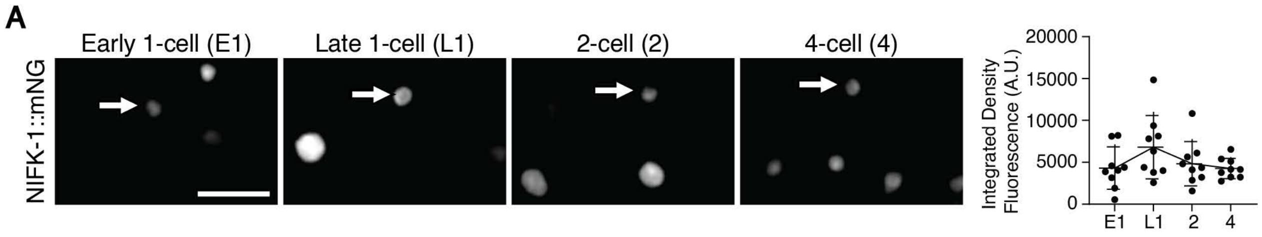
A
Early 1-cell (E1)
Late 1-cell (L1)
2-cell (2)
4-cell (4)
NIFK-1::mNG
Integrated Density Fluorescence (A.U.)
20000
15000
10000
5000
0
E1
L1
2
4

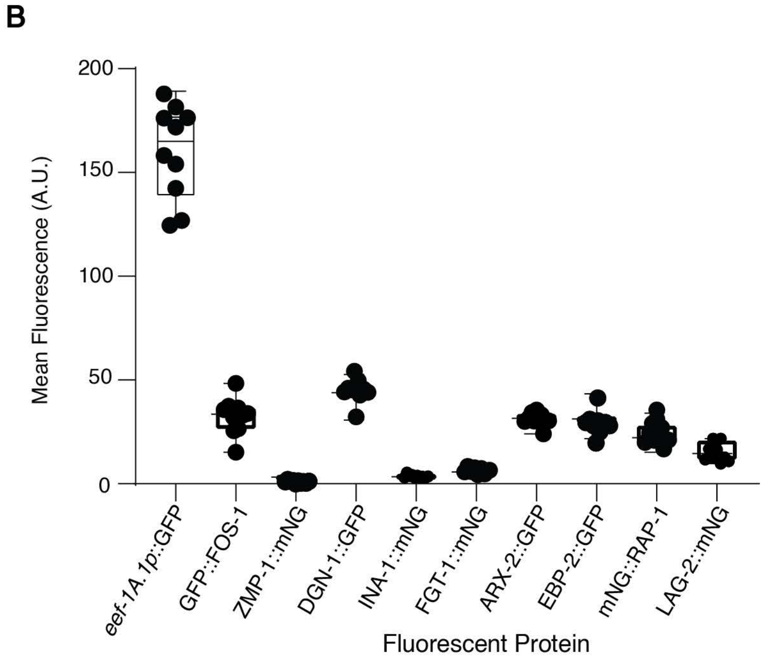
B
Mean Fluorescence (A.U.)
200
150
100
50
0
eef-1A.1p::GFP
GFP::FOS-1
ZMP-1::mNG
DGN-1::GFP
INA-1::mNG
FGT-1::mNG
ARX-2::GFP
EBP-2::GFP
mNG::RAP-1
LAG-2::mNG
Fluorescent Protein

**Figure S5. Ribosome biogenesis protein NIFK-1 spikes in levels at the P6.p late 1-cell stage and the *eef-1a.1* promoter is expressed robustly.**

(A) The timeline of endogenous NIFK-1 (NIFK-1::mNG) expression in the AC (arrow). NIFK-1 shows a spike in levels at the P6.p late 1-cell stage. Graph shows quantification of total fluorescence levels (right) over developmental time ($n \geq 10$ ACs per timepoint, mean ± SD, one-way ANOVA with post hoc Tukey's test). Quantification is from one experiment in which several animals at each timepoint were processed in parallel. Scale bar: 5 μm.

(B) Fluorescence intensity of *eef-1a.1p*::GFP at the P6.p 2-cell stage is higher than all endogenously-tagged pro-invasive proteins ($n \geq 8$ ACs per strain, boxplot).
